# Fatty Acid Synthase inhibitor TVB-3166 prevents S-acylation of the Spike protein of human coronaviruses

**DOI:** 10.1101/2020.12.20.423603

**Authors:** Katrina Mekhail, Minhyoung Lee, Michael Sugiyama, Audrey Astori, Jonathan St-Germain, Elyse Latreille, Negar Khosraviani, Kuiru Wei, Zhijie Li, James Rini, Warren L. Lee, Costin Antonescu, Brian Raught, Gregory D. Fairn

**Affiliations:** Department of Biochemistry, University of Toronto, Toronto, Ontario, Canada; Keenan Research Centre, St. Michael’s Hospital, Unity Health Toronto, Toronto, Ontario, Canada; Department of Chemistry and Biology, Ryerson University, Toronto, Ontario, Canada; Princess Margaret Cancer Centre, University Health Network, Ontario, Canada; Department of Medical Biophysics, University of Toronto, Ontario, Canada; Department of Laboratory Medicine and Pathobiology, University of Toronto, Toronto, Ontario, Canada; Department of Molecular Genetics, University of Toronto, Ontario, Canada; Department of Medicine, University of Toronto, Toronto, Ontario, Canada; Department of Surgery, University of Toronto, Toronto, Ontario, Canada; Department of Pathology, Dalhousie University, Halifax, Nova Scotia, Canada

**Keywords:** S-acylation, ZDHHC, Spike coronavirus, click chemistry, fatty acid synthase

## Abstract

The Spike protein of SARS-CoV2 and other coronaviruses mediate host cell entry and are S-acylated on multiple phylogenetically conserved cysteine residues. Multiple protein acyltransferase enzymes of the ZDHHC family have been reported to modify Spike proteins post-translationally. Using resin-assisted capture mass spectrometry, we demonstrate that the Spike protein is S-acylated in SARS-CoV2 infected human and monkey cells. We further show that increased abundance of the human acyltransferase ZDHHC5 results in increased S-acylation of the SARS-CoV2 Spike protein, whereas *ZDHHC5* knockout cells had a 40% reduction in the incorporation of an alkynyl-palmitate using click chemistry detection. We also find that the S-acylation of the Spike protein is not limited to palmitate, as clickable versions of myristate and stearate were also found on the immunocaptured protein. Yet, ZDHHC5 was highly selective for palmitate, suggesting that other ZDHHC enzymes mediated the incorporation of other fatty acyl chains. Thus, since multiple ZDHHC isoforms may modify the Spike protein, we examined the ability of the fatty acid synthase inhibitor TVB-3166 to prevent the S-acylation of the Spike proteins of SARS-CoV-2 and human CoV-229E. Treating cells with TVB-3166 inhibited S-acylation of ectopically expressed SARS-CoV2 Spike and attenuated the ability of SARS-CoV2 and human CoV-229E to spread *in vitro.* Additionally, treatment of mice with a comparatively low dose of TVB-3166 promoted survival from an otherwise fatal murine coronavirus infection. Our findings further substantiate the necessity of CoV Spike protein S-acylation and the potential use of fatty acid synthase inhibitors.

## Introduction

Since 2002, coronaviruses (CoV) have resulted in three zoonotic outbreaks: SARS in 2002, MERS in 2012, and SARS-CoV2 in December 2019 (1, 2). CoVs infect humans and animals and cause various diseases targeting different tissues, including respiratory, enteric, renal, and hepatic (3, 4). The current pandemic highlights the need for effective treatments of this family of viruses. There are enormous efforts to develop prophylaxis and therapeutic options for SARS-CoV2, including vaccination, protease inhibitors, and soluble decoy receptors (5–7). Many putative countermeasures target the Spike protein, which is required to attach the virus to cells via binding to the angiotensin-converting enzyme (ACE2) and potentially other host proteins (6). Further, Spike protein proteolytic cleavage generates an S2 fragment capable of stimulating viral fusion and payload delivery into the cytosol (8). However, the high mutation rates of positive-sense (+) sense RNA viruses may result in resistance to some of these therapeutic approaches (9, 10). Additionally, many potential zoonotic coronavirus species in bat and civet populations necessitate exploring all available therapies (11, 12). Thus, identifying a pan-CoV infection therapy that targets a host enzyme is desirable for current and future potential zoonotic infections.

Although not truly synonymous, S-palmitoylation or S-acylation are often used interchangeably to describe the reversible covalent addition of palmitoyl or other fatty acyl chains to cysteine (Cys) residues via a thioester bond. This modification can occur spontaneously in high concentrations of acyl-CoAs or catalyzed by a family of zinc finger Asp-His-His-Cys-containing protein acyltransferases or simply ZDHHC enzymes. In the Spring of 2020, Krogan and colleagues reported that ZDHHC5 and its binding partner Golga7 interact with the Spike protein of SARS-CoV2 using an affinity purification/mass spectrometry approach (13). The results suggest that the cytosolic tail of the Spike protein may be post-translationally S-acylated. A variety of labs have substantiated this, and collectively, these studies have demonstrated that multiple protein acyltransferases, including ZDHHC2, 3, 5, 6, 8, 9, 11, 12, 20, 21 and 24, directly or indirectly influence the S-palmitoylation/acylation of the Spike protein depending on the cellular context (14–20). These findings likely result from the lack of specificity of some ZDHHC enzymes and the promiscuous nature of some substrates. Considering that the Spike protein contains ten Cys residues in proximity to the membrane, this would only increase the chances of modification by various transferases.

Previously, the Spike protein of the murine hepatitis virus (MHV) required S-acylation of its C-terminal tail to support virion production (21). Specifically, preventing S-acylation prevented the MHV Spike protein from being incorporated into new virions (21). Further, a mutant version of Spike unable to be S-palmitoylated displayed a reduced ability to stimulate membrane fusion vis-à-vis syncytium formation field (21, 22). Additionally, S-acylation of the SARS Spike is critical for its ability to catalyze cell-cell fusion (23). Here we aimed to extend the findings and investigate the role of ZDHHC5 in Spike S-acylation and its role in the human cold virus CoV-229E. Given the large number of ZDHHC enzymes that may potentially S-acylate the Spike protein, we also sought to examine the ability of the fatty acid synthase inhibitor TVB-3166 to limit S-acylation.

## Results

### SARS-CoV2 Spike protein is S-acylated

Alignment of the C-terminal cytoplasmic tails of several human and murine CoVs reveals that the 20 amino acids adjacent to the membrane are highly enriched in Cys residues (Fig. 1A). Indeed, in the case of SARS-CoV2 Spike, half of the first 20 amino acids are Cys residues, providing ample sites for potential S-acylation. To explore this possibility, we first established a model to express and study the SARS-CoV2 Spike protein using ectopic expression of Spike tagged at the C-terminus with a C9 epitope (TETSQVAPA) (Fig. 2A) (24, 25). Immunoblotting of the Spike-C9 revealed numerous reactive bands in transfected cells but none in controls (Fig. 2B). Consistent with previous reports (25, 26), we detected several Spike molecular species, including a full-length ∼190 kDa and a ∼100 kDa fragment compatible with the furin-cleaved S2 fragment. To determine if the ectopically expressed Spike-C9 protein is acylated, we incubated transfected cells with BSA conjugated alkynyl-palmitate (15-hexadecynoic acid; 15-HDYA), a “clickable” palmitate analog (27). Spike-C9 immunocaptured using anti-C9 beads was reacted with cyanine 5.5 (Cy5.5) – azide using a copper(I)-catalyzed azide-alkyne cycloaddition (CuAAC) (28–30). As shown in Fig. 1B, the Spike-C9 from cells incubated with alkynyl-palmitate and reacted with the Cy5.5-azide generated a similar banding pattern as the anti-C9 antibody, indicating that the full-length and S2 fragment of the Spike protein are both acylated.

**Figure 1.**
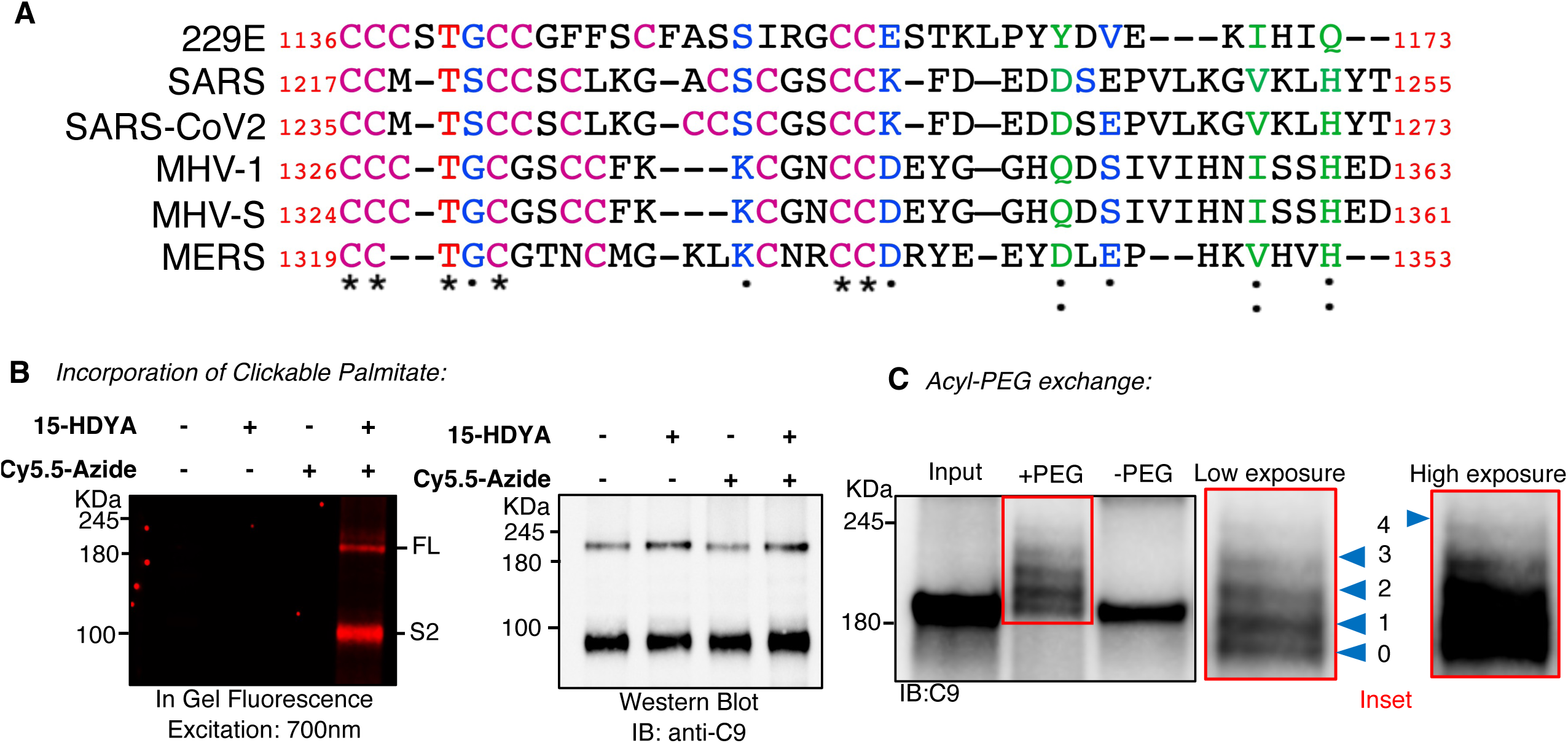
S-Acylation of the SARS-CoV2 Spike protein. **(A)** Sequence alignment of the cytosolic amino acids adjacent to the transmembrane domain of murine and human coronavirus Spike proteins. Adapted from ClustalW; (*) residue is wholly conserved, (:) strongly similar properties, (.) weakly similar properties conserved. **(B)** Incorporation of 15-HDYA (ω-terminal alkyne containing palmitate analog) on Spike-C9 measured through click chemistry. The covalent addition of 15-HDYA to the Spike-C9 is determined following immunocapture using anti-C9 beads and a copper-catalyzed cycloaddition between the alkyne group Cy5.5-conjugated Azide. Samples were resolved using gel electrophoresis and imaged using an LI-COR Odyssey (depicted as in-gel fluorescence and the left). Subsequently, samples were transferred to PVDF membrane for immunoblotting with an anti-C9 antibody followed by an HRP-conjugated secondary antibody and chemiluminescence (western blot on the right). Omission of either the 15-HDYA or the Cy5.5-Azide results in loss of specific fluorescence. N =3 biological replicates **(C)** Examination of the number of putative S-acylation sites. Lysates of HEK293T cells expressing the Spike-C9 epitope were subjected to the Acyl – polyethylene glycol (PEG) exchange assay. Spike-C9 was immunocaptured using anti-C9 beads followed by the blocking of free thiol groups, cleavage of the thioester bond with hydroxylamine and the covalent addition of the 5 kDa PEG to the freshly liberated thiol residues. Following the acyl-PEG assay samples were resolved by electrophoresis and transferred to membranes for blocking and detection. Input: unprocessed HEK293T lysate, +PEG: de-acylated and PEG covalent addition, and -PEG: de-acylated without PEG addition. The inset of the acyl-PEG assay immunoblot (on the right) at low and high exposures demonstrates an unmodified protein and 4 addition bands indicating a reaction of the Spike has at least 4 sites of S-acylation, n=3 biological replicates.

**Figure 2.**
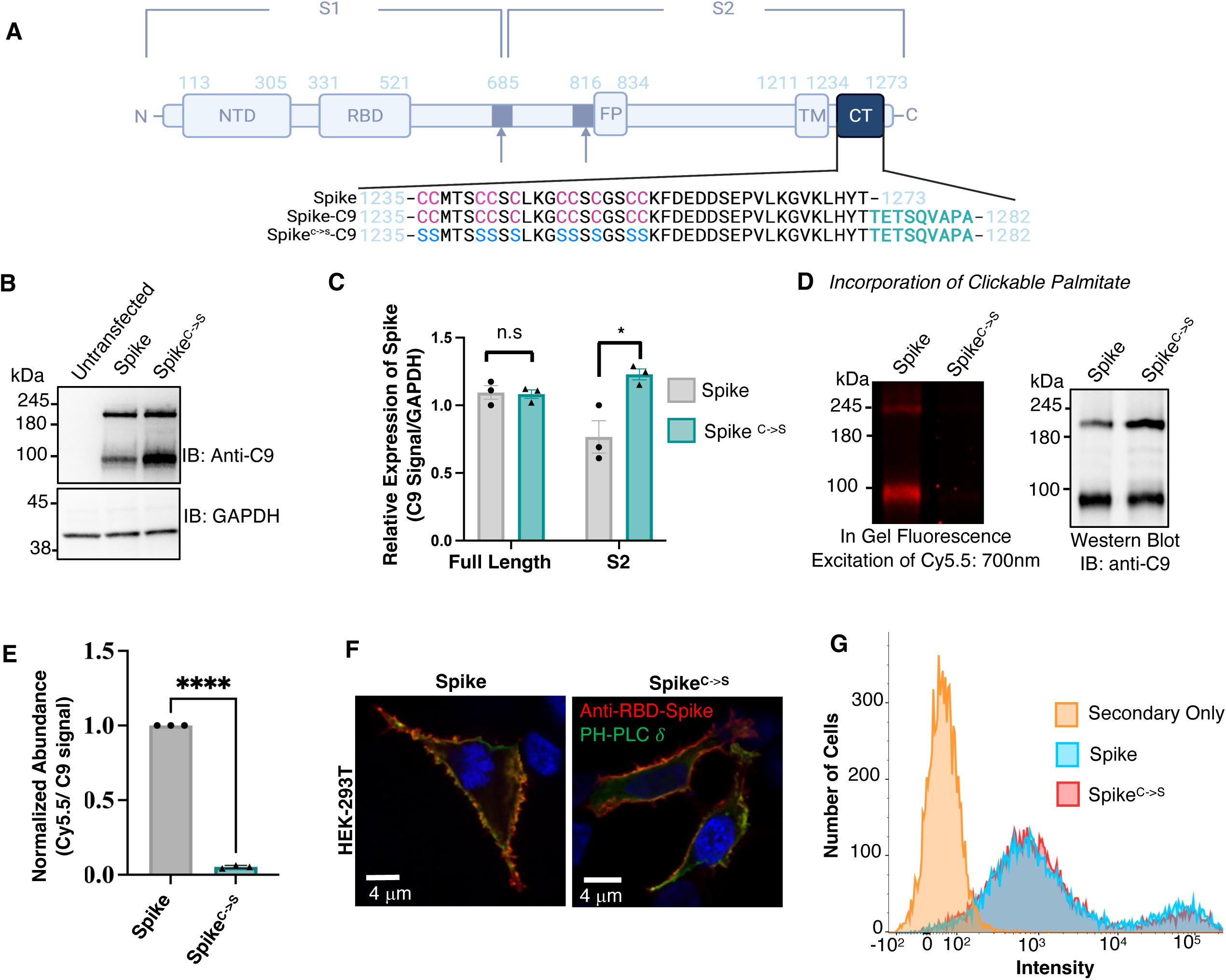
Cytosolic Cysteine Residues are required for the addition of clickable palmitate but not transit to the cell surface. **(A)** Schematic representation of the SARS-CoV2 Spike protein and amino acid sequence of the cytosolic amino acids adjacent to the transmembrane domain, the all cysteine (Magenta) to serine (Cyan) substitution and the location of the C9 epitope in relation to the sites of protein lipidation. Arrows depict cleavage sites, NTD: N-terminal domain, RBD: receptor binding domain, FP: fusion peptide, CT: C-terminus. **(B)** Immunoblots of ectopically expressed Spike-C9 and Spike^C->S^ -C9 and GAPDH from total cell lysates of HEK293T cells. Cells were transiently transfected for 18h prior to collection and subsequent immunoblotting with the anti-C9 antibody. Untransfected cells display no anti-C9 reactive bands whereas cells transfected with Spike-C9, or the all Cys-to-Ser mutant have bands equivalent to the full-length protein and a cleaved S2 fragment. **(C)** Quantitation of panel ‘B’. Data are means ± SEM of n=3 biological replicates. Unpaired t-test with Welch’s corrections, ns = not significant p = 0.85 and * p = 0.02. **(D)** Clickable palmitate is not covalently attached to Spike-^C->S^. Cell expressing either Spike-C9 or Spike^C->S^-C9 were incubated with 15-HDYA and subsequently processed using copper-catalyzed cycloaddition of Cy5.5-Azide as in Figure 1B. The degree of protein lipidation was detected by in-gel fluorescence of Cy5.5-Azide (exposed at 700 nm; left of the image); corresponding protein levels were detected by chemiluminescence (right of the image). **(E)** Quantitation of panel ‘D’ and the labelling by exogenously added 15-HYDA. The abundance is presented as the quotient of the intensities of the Cy5.5 signal from the attached fatty acid and the C9 epitope detected by immunoblotting. Data are means ± SEM of n=3 biological replicates and analyzed using an unpaired t-test with Welch’s correction; **** P<0.0001. **(F)** Confocal imaging of surface-exposed and immunostained Spike and Spike^C->S^. Immunofluorescence of Spike-C9 and Spike^C->S^-C9 ectopically expressed and stained with an anti-Receptor Binding Domain (RBD) of Spike antibody (red) in non-permeabilized HEK293T cells. Cells were co-transfected with the pleckstrin homology domain of phospholipase Cδ (GFP-PH-PLCδ), which delineated the plasma membrane, n=3 biological replicates. **(G)** Flow cytometry of surface-exposed Spike and the S-acylation deficient mutant as in ‘F’, n=3 biological replicates.

Considering that the SARS-CoV2 Spike protein cytoplasmic tail contains ten cysteine residues, we sought to confirm that the modifications are on Cys residues and the degree of modification. To do this, we used an approach termed <u>a</u>cyl-<u>p</u>olyethylene glycol (PEG) <u>e</u>xchange (APE) that involves substituting acyl groups attached to Cys residues with a 5-kDa PEG mass tag field (31). The unreacted protein (input) and the protein treated with hydroxylamine to cleave the thioester bond and remove the fatty acyl chains, but not reacted with PEG (-PEG), have an apparent molecular weight of ∼190 kDa (Fig. 1C). Spike-C9 treated with hydroxylamine and reacted with a maleimide-functionalized five kDa PEG revealed five distinct bands consistent with an unmodified band and four sites of S-acylation. Given the proximity of the ten Cys residues within a 20 amino acid stretch, it is unclear if the addition of multiple five kDa PEG molecules could be sterically hindered. Another possibility is that the addition of the C9 epitope to the C-terminal end of the Spike may result in less or variable acylation. Regardless, these results demonstrate that the Spike protein can be modified on numerous sites and that some individual proteins are modified with a minimum of four acyl chains.

Next, we sought to confirm the results obtained using the heterologously expressed epitope-tagged Spike with the native Spike in cells infected with the SARS-CoV2 virus. We used a strategy termed acyl resin-assisted capture (acyl-RAC) (32). In this workflow, free Cys residues are chemically blocked using N-ethylmaleimide. Next acyl-cysteine thioester linkages are then cleaved using hydroxylamine, and then treated lysates are incubated with thiol-reactive beads allowing for the capture of the previously S-acylated proteins by a mixed disulphide exchange reaction. This assay is readily coupled to mass spectrometry (MS) based protein detection and quantitation. As shown in Table 1, Table S1, and S2, the Acyl-RAC MS workflow can identify numerous host proteins that are known to be S-acylated, including flotillin, calnexin, scribble and SNAP23 in both Vero and HEK293 stably expressing both the ACE2 receptor and TMPRSS2 protease (or simply HEK A2T2) (33). Analysis of cells infected with SARS-CoV2 revealed the presence of the Spike protein and that it was one of the most abundant S-acylated proteins in both cell types. Furthermore, despite the infection, most of the other host S-acylated proteins were unaltered. These results collectively demonstrate that both the ectopically expressed Spike and Spike delivered during SARS-CoV2 infection are S-acylated.

**Table 1.** Most abundant S-acylated proteins in control and SARS-CoV2 infected Vero and HEK cells expressing Ace2 and TMPRRS2. Proteins identified using Acyl Resin Assisted Capture (Acyl-RAC) of control and SARS-CoV2 infected Vero and HEK A2T2 cells. Hydroxylamine is used to distinguish S-acylated proteins, whereas NaCl treated samples are used for non-specific binding as described in the Experimental Procedures. Full lists are available in the Supplemental Tables.

### S-acylation of SARS-CoV2 Spike protein is required for membrane fusion

The CoV Spike polypeptides are class I viral fusion proteins that mediate membrane fusion (34). This function is essential to initiating viral infection. To investigate the role of S-acylation of the SARS-CoV-2 Spike protein in this process, we used a site-directed mutagenesis approach to mutate all ten Cys residues in the cytoplasmic tail to Ser (termed Spike^C->S^) (Fig 2A). Lysates of HEK293T cells ectopically expressing Spike-C9 and Spike^C->S^-C9 were immunoblotted and revealed that the full-length Spike^C->S^-C9 was expressed at comparable levels to the wild-type protein and that there is also the production of the S2 fragment (Fig. 2B, C). Our studies noticed that the relative proportion of full-length compared to the S2 was variable but consistent within individual biological replicates. Whether this is due to a cell-autonomous factor, the efficiency of transfection, or perhaps even post cell lysis proteolysis was not determined. We next confirmed that the Spike^C->S^-C9 is not S-acylated by measuring the addition of the 15-HDYA (Fig 2D, E). Recent studies have demonstrated that the C-terminal region of Spike interacts with a variety of vesicular transport machinery, including COPI and COPII coatomers (35). To determine if loss of the S-acylation sites alters the protein delivery to the plasma membrane, we used an antibody that detects an extracellular epitope together with immunofluorescence microscopy and flow cytometry. We confirmed that comparable amounts of both the Spike-C9 and Spike^C->S^-C9 traffic to the plasma membrane (Fig. 2F, G). Together these results demonstrate that mutagenesis of the cytoplasmic Cys residues to Ser and the concomitant loss of S-acylation have minimal impact on expression levels and transport to the cell surface.

Previous studies have demonstrated that the SARS-CoV-2 Spike protein can catalyze syncytium formation, provided that neighbouring cells express ACE2 (6). To determine if S-acylation of the Spike protein is required for syncytium formation, we utilized a co-culture strategy where cells co-transfected with plasmids encoding Spike-C9 and soluble mCherry were co-cultured with cells co-transfected with plasmids encoding ACE2 and soluble GFP. Co-cultures containing cells expressing mCherry alone and cells with ACE2/GFP showed no signs of cell-cell fusion, as expected (Fig. 3A, B). However, co-cultures with Spike/mCherry and ACE2/GFP displayed numerous large syncytia containing soluble GFP and mCherry (Fig. 3 A, B). In contrast, HEK293T cells expressing Spike^C->S^-C9/mCherry co-cultured with ACE2/GFP expressing cells displayed only a few smaller mCherry and GFP double-positive cells (Fig 3A, B). These results demonstrate that one or more membrane-proximal cytoplasmic Cys residues are required for the Spike protein to facilitate syncytium formation.

**Figure 3.**
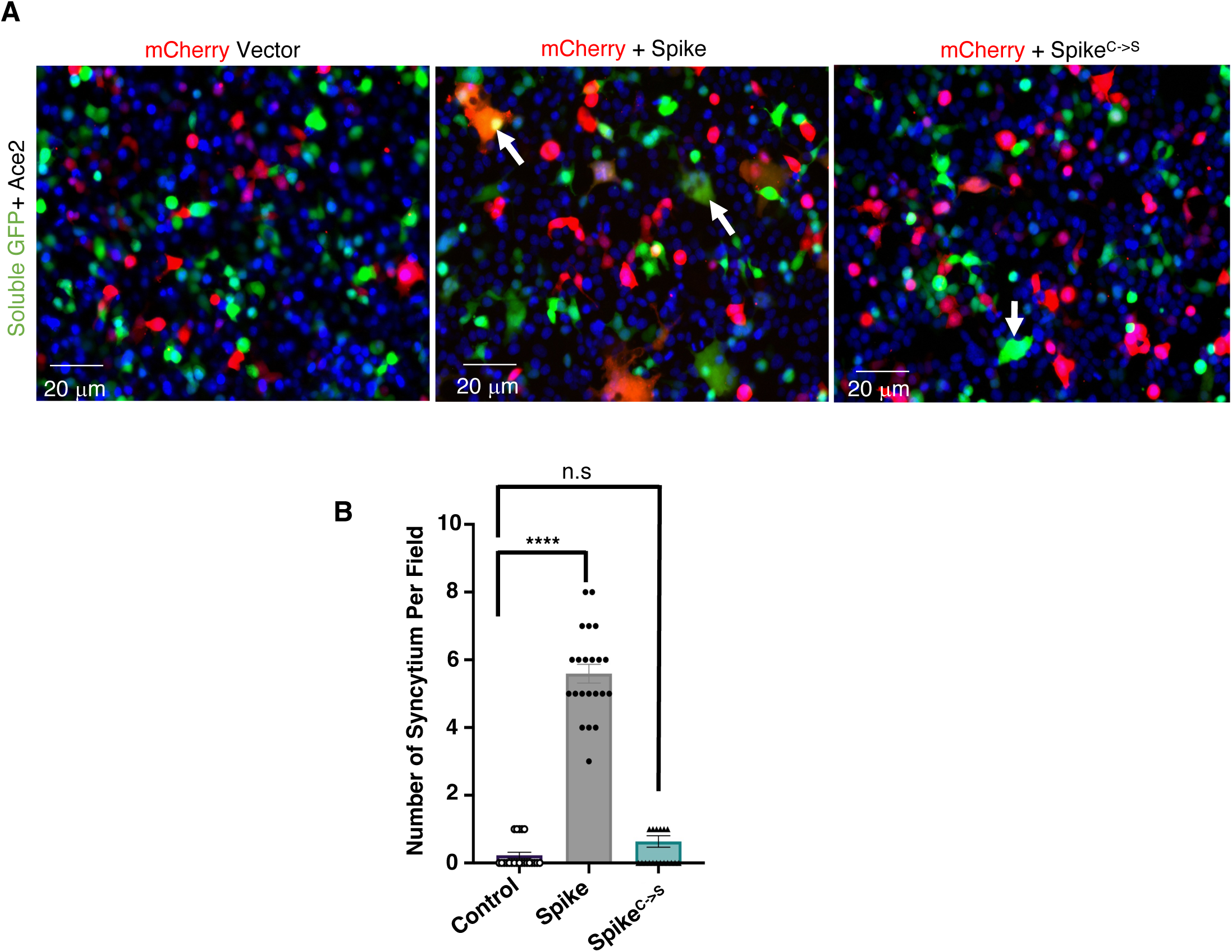
S-acylation of the SARS-CoV2 Spike protein is required for cell-cell fusion. **(A)** Confocal images of mCherry and GFP expressing co-culture to assess syncytium formation. HEK293T cells expressing mCherry and empty vector, Spike-C9 or Spike^C->S^-C9 were plated with HEK293T cells expressing soluble GFP and the ACE2 receptor. Representative two-color merged micrographs demonstrate that wild-type Spike and ACE2 co-culture form syncytium (Orange) indicated with white arrows. **(B)** Quantitation of panel ‘A’, the number of syncytia per microscope field of view were counted, and 10 microscope fields of each condition per experiment were analyzed. Data are means ± SEM of n= 3 biological replicates and analyzed by one-way ANOVA; **** P<0.0001, n.s P=0.2418 compared to control.

### ZDHHC5 is required for human coronavirus infection

The human genome encodes a family of 24 ZDHHC (including ZDHHC11B) palmitoyltransferases (*22*). ZHHC5 was reported to interact with the SARS-CoV2 Spike protein physically, as shown using IP-MS (13). We confirmed these results using co-immunoprecipitation and found that Spike-C9 could transiently interact with HA-tagged ZDHHC5 (Fig. S1). Given that numerous ZDHHC enzymes have been demonstrated to S-acylate the Spike protein, we wanted to determine the relative importance of ZDHHC5. Thus, we assessed the contribution of ZDHHC5 to the S-acylation of Spike in parental HEK293T cells and genome-edited cells deficient in ZDHHC5 expression (Fig. S2). Spike-C9 was transiently expressed in wild-type and ZDHHC5^ko^ cells, incubated with 15-HDYA and subsequently processed as in Figure 1 to determine the extent of S-palmitoylation. Notably, the loss of ZDHHC5 resulted in nearly a 40% reduction in Spike S-palmitoylation (Fig 4A, B). Thus, even though reports demonstrate that upwards of 11 ZDHHC enzymes can modify Spike, our experiments with the clickable palmitate suggest ZDHHC5 is responsible for a significant fraction of the post-translation modification.

**Figure 4.**
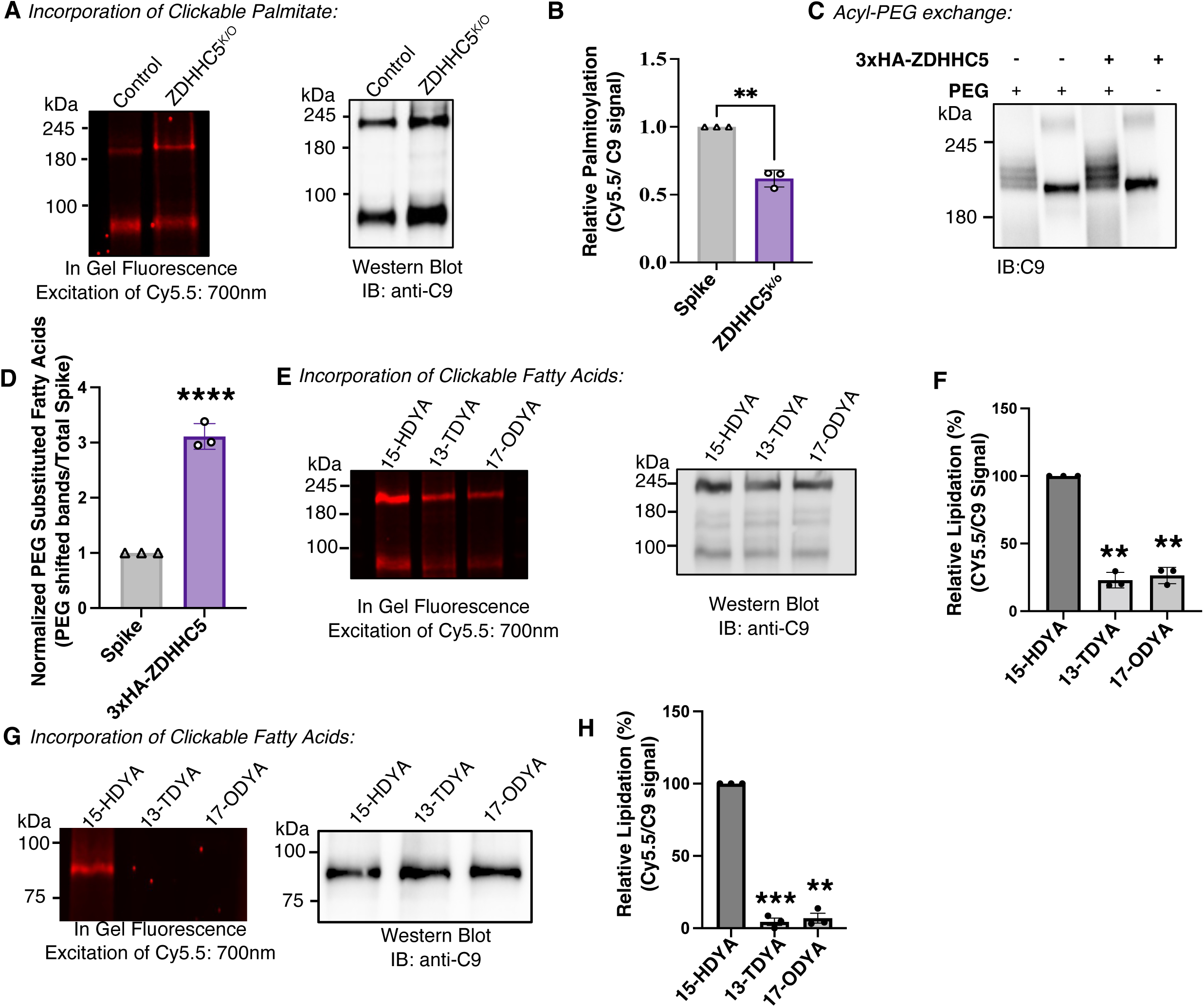
The acyltransferase ZDHHC5 contributes to the S-acylation of Spike with a preference for palmitate. **(A)** Control and genome-edited *ZDHHC5* knockout HEK293T cells were incubated with 15-HDYA and processed using a click chemistry reaction (Figure 1B). Covalent attachment of the clickable palmitate analog was determined by Cy5.5 in-gel fluorescence (exposure 700 nm; left of the image), with the corresponding protein levels determined following transfer to membrane and immunoblotting using chemiluminescence (right of the image). **(B)** Quantitation of panel ‘A’. Relative palmitoylation levels were calculated as the quotient of the Cy5.5 signal and the C9 chemiluminescence. Data are the means ± SEM of n=3 biological replicates and analyzed using an unpaired t-test with Welch’s correction. ** P=0.0089. (**C)** Control cells and cells transiently transfected with 3xHA-ZDHHC5 were processed using the Acyl-PEG exchange assay as in Figure 1C. Following the 5-kDa PEG substitution, samples were resolved, transferred, and immunoblotted using the anti-C9 antibody. **(D)** Quantitation of panel ‘C’ whereby the shifted PEGylated bands of the 180 kDa Spike are normalized to the unmodified band at ∼180 kDa; this band migrated to the same position as the minus PEG sample. In individual experiments, the value of the control cells was set equal to 1, and the relative intensity of the ZDHHC5 over-expressing cells was determined as a relative measurement. Data is the mean ± SEM of n=3 biological replicates and analyzed with an unpaired t-test with Welch’s correction; **** P<0.0001 compared to controls. **(E)**. Cell co-expressing Spike-C9 and 3xHA-ZDHHC5 were incubated with alkyne containing clickable analogs of myristate (13-TDYA) and palmitate (15-HDYA) and stearate (17-ODYA) for 2 hours. The spike protein was immunocaptured and processed as in Figure 1B. In-gel fluorescence of the Cy5.5 (left) and immunoblots of the anti-C9 (right) are shown. **(F)** Relative lipidation of Spike-C9 whereby the Cy5.5-azide in-gel fluorescence was normalized to the total Spike-C9 protein levels measured by chemiluminescence. Data are the mean ± SEM of n=3 individual trial replicates analyzed by multiple t-tests with Welch’s correction; ** P<0.001 compared to 15-HDYA. **(G)** Cells ectopically expressing 3xHA-ZDHHC5 were incubated with clickable fatty acids as in panel ‘E’. Immunocapture using an anti-HA antibody and Protein A beads were used to purify 3xHA-ZDHHC5 from lysates and subjected to click chemistry analysis as in panel ‘E’. The results demonstrate that ZDHHC5 cannot undergo auto-acylation with the myristate and stearate analogs. **(H)** Relative lipidation of 3xHA-ZDHHC5 whereby the Cy5.5-azide in-gel fluorescence was normalized to the total 3xHA-ZDHHC5 protein levels measured by chemiluminescence. Data are the mean ± SEM of n=3 individual trial replicates analyzed by multiple t-tests with Welch’s correction; *** P=0.0007 ** P<0.001 compared to 15-HDYA.

As mentioned previously, the 15-HDYA experiment doesn’t provide information about the number of sites being modified. Furthermore, 15-HDYA does not provide information on whether other fatty acids can be attached to the Spike protein. We sought to investigate this using complementary approaches. First, cells ectopically expressing Spike-C9 and 3xHA-ZDHHC5 were processed using the APE assay as in Figure 1C. The increased expression of ZDHHC5 resulted in a ∼3-fold increase in S-acylation as detected by APE, along with the appearance of a 6^th^ band at a higher molecular weight (Fig 4C, D).

Unlike the metabolic labelling with clickable fatty acids, the APE assay detects sites of S-acylation regardless of the source of the acyl chains. Next, we considered whether the Spike protein was modified by acyl chains other than palmitate and whether over-expression of the ZDHHC5 enhanced this modification. To investigate this possibility, cells were metabolically labelled with BSA-conjugated clickable fatty acid analogs of palmitate, stearate and myristate, followed by immunocapture and processing as in Figure 1B. As depicted in Fig 4E, F, the myristate analog (13-TDYA) and stearate analog (17-ODYA) are covalently attached to the Spike-C9 but less abundant. Curiously we also found that over-expression of ZDHHC5 had a modest impact on the labelling of Spike by 15-HDYA compared to control cells (Fig S3). We hypothesize that this is likely because either the uptake from the medium and synthesis of the 15-HDYA-CoA are limiting steps, and in the presence of *de novo* synthesized palmitoyl-CoA, it is inefficiently used to S-acylate proteins. Indeed when we repeat the experiment in the presence of the fatty acid synthase inhibitor TVB-3166, we see more efficient incorporation of the clickable label (Fig S3).

As part of the reaction cycle, ZDHHC enzymes are first auto-acylated with the acyl chains and subsequently transferred to the substrates (36). To our knowledge, no information is available on the fatty acyl specificity of ZDHHC5. Thus, taking advantage of the clickable fatty acids and protocol used in Fig 4E, we sought to determine which fatty acids are attached to ZDHHC5 as part of the reaction cycle. As shown in Fig 4G and H, ZDHHC5 displays a strong preference for 16-carbon acyl chains even compared to the 14- and 18-carbon chains. The results suggest that ZDHHC5 is responsible for a substantial amount of the palmitate attached to the Spike protein, but that other ZDHHC enzymes attach other fatty acyl chains to the Spike protein.

### ZDHHC5 supports human CoV-229E Spike S-acylation and infection

Next, we sought to investigate if our observations on the Spike protein and ZDHHC5 can be extended to another human coronavirus, namely CoV-229E, one of the viruses that cause the common cold. Although the Spike protein of CoV-229E recognizes a different host protein, CD13 (37), its cytosolic tail is similar to that of SARS-CoV2 (Fig. 1A). We again generated a plasmid-borne C9 tagged Spike protein of the 229E virus (229E Spike-C9). Using the APE assay, we found that the 229E Spike-C9 protein is S-acylated on multiple sites when ectopically expressed in HEK293T cells. Curiously, the pattern obtained was different from that of the SARS-CoV2 Spike. First, we found that all the 229E Spike-C9 had at least one PEG attached (Fig 5A) and that we could not resolve the individual bands. This raises the possibility that the 229E Spike is more heavily S-acylated than the SARS-CoV2 Spike. Again, the intensity of the PEGylated proteins is enhanced when HA-ZDHHC5 levels are increased (Fig. 5A, B). This finding is consistent with the Spike proteins of SARS-CoV2 and likely 229E being substrates of ZDHHC5. To our knowledge, the role of other ZDHHC enzymes and the 229E Spike has not been investigated. Our results do not rule out the possibility that other ZDHHC enzymes may also modify the 229E Spike protein.

**Figure 5.**
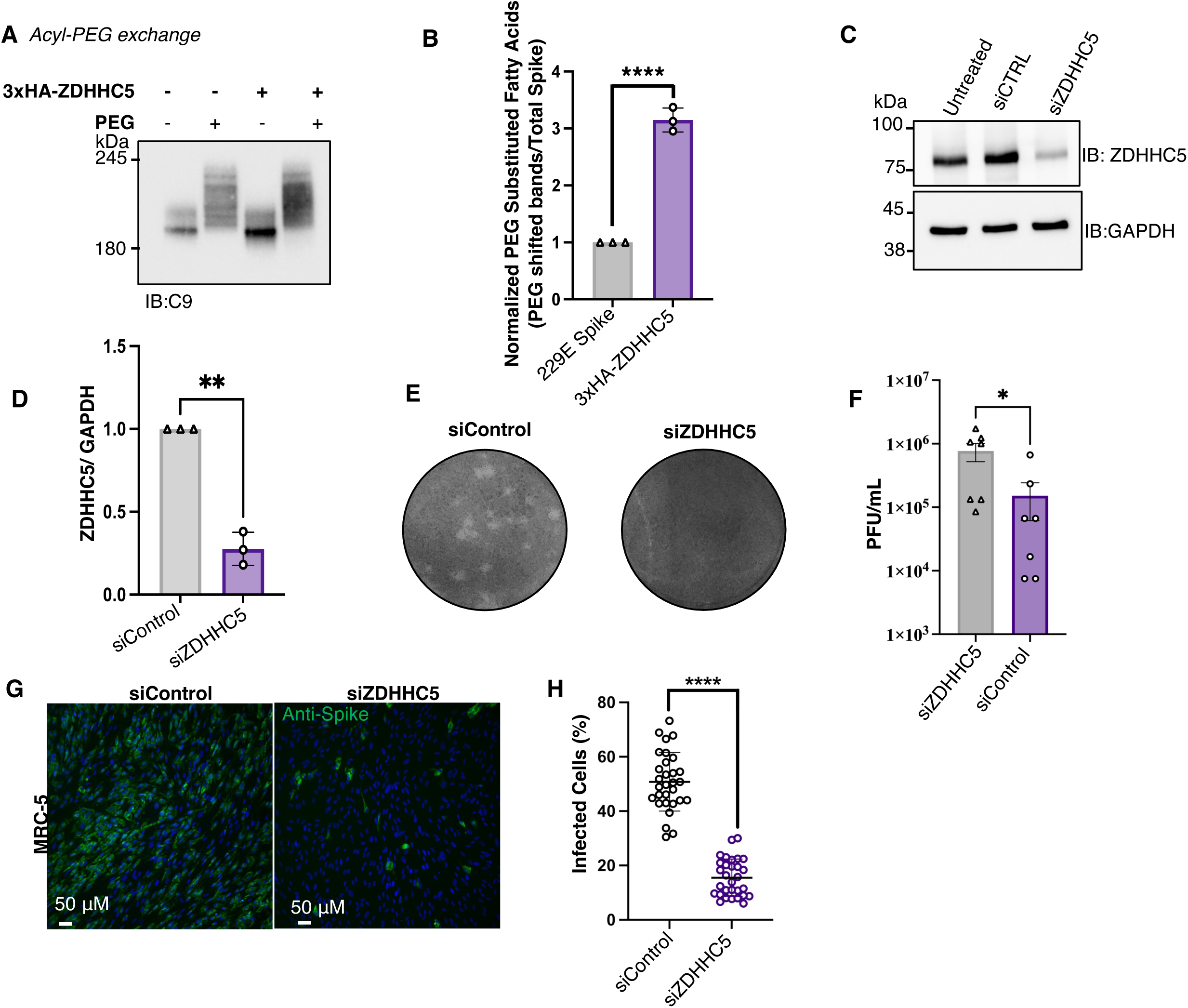
Increased expression of *ZDHHC5* increases S-acylation of human CoV 229E Spike and is required for virulence. **(A)** Acyl-PEG assay as in Figure 1C to examine the S-acylation of the 229E Spike-C9 in HEK293T cells transiently transfected with either the 3xHA-ZDHHC5 or empty vector. **(B)** Quantitation of the acyl-PEG assay in panel ‘A’ where the higher molecular weight PEG-modified bands were normalized to the fastest migrating band. All bands in the +PEG sample migrated more slowly than the -PEG sample suggestion the 229E Spike is more readily and perhaps more highly S-acylated. Data are the mean ± SEM of n=3 biological replicates, unpaired t-test with Welch’s correction, **** P<0.0001. **(C)** MRC-5 cells were transfected with siRNA-control or siRNA-ZDHHC5, and lysates were collected and immunoblotted to examine the relative expression of ZDHHC5. **(D)** Quantification of panel ‘C’. Data are the mean ± SEM n = 3 biological replicates normalized to GAPDH and analyzed using an unpaired t-test with Welch’s correction, **P=0.0063. **(E)** Representative image of plaques observed in MRC-5 cells silenced for ZDHHC5 infected with 229E for 5 days. **(F)** Quantification of 229E PFU/mL in MRC-5 cells silenced for ZDHHC5 as shown in panel ‘E’, unpaired t-test, * P=0.0371. Data are mean ± SEM of n=3 biological replicates **(E)** Representative images of MRC-5 cells silenced for ZDHHC5 and infected with 229E at an MOI of 0.0005. Images labelled as follows: Spike 229E (green), Dapi (blue) (**F**) Quantification of 229E infection assay where each data point represents a 10x magnified field of view where an average of 600 cells were counted in ImageJ, 10 fields of view per condition for n=3 biological replicates were analyzed; unpaired t-test, ****p< 0.0001. Data are mean ± SEM.

To investigate the role of ZDHHC5 in the generation of virulent progeny, we used two complementary cell-based assays. First, we used a previously described siRNA to silence *ZDHHC5* in confluent monolayers of human MRC-5 fibroblasts (Fig. 5C, D) (38). Next, we used these conditions and assessed the ability of CoV-229E to form plaques (Fig. 5E, F). Cell monolayers transfected with a non-targeting siControl produced an average of ∼10^6^ PFU/ml, and monolayers transfected with the siZDHHC5 displayed significantly fewer plaques. From the plaque assay, it was unclear if the loss of ZDHHC5 impacted the initial infection or the subsequent spread to the neighbouring cells in the monolayer. We used a liquid culture assay to complement the plaque assay to investigate infection and dissemination. In this assay, cells were incubated in the presence of 229E virus at a low multiplicity of infection (MOI = 0.005) for 2 hours, followed by extensive washing. After 72 hours, cells were processed and stained with an antibody directed against the 229E Spike protein (39) and an AlexaFluor 488-conjugated secondary antibody. As shown in Fig. 5G and H, ≈50% of the siControl cells were infected as determined by immunofluorescence detection of the Spike protein. In contrast, only ≈15% of the ZDHHC5 silenced cells were positive for the Spike protein. These results are consistent with the notion that ZDHHC5 is required for the generation of virulent progeny and spread in vitro.

### *De novo* fatty acid synthesis is required for CoV infectivity

Based on our data, we sought to determine if ZDHHC5 and other ZDHHC enzymes could represent suitable therapeutic targets for treating CoV infections. Unfortunately, no specific inhibitor of ZDHHC5 exists. Previously, we found that the fatty acid synthase (FASN) inhibitor cerulenin prevented the S-palmitoylation of the ZDHHC5 substrates NOD1 and NOD2 (40). This is logical since FASN produces a cytosolic pool of palmitoyl-CoA, the palmitate donor for ZDHHCs. However, cerulenin is not suitable therapeutically due to severe weight loss and off-target effects in murine models (41–43). Fortunately, first-in-class FASN inhibitors have been developed by Sagimet Biosciences, formerly 3-V Biosciences (San Mateo, USA). One of these inhibitors, TVB-2640, is orally available, well-tolerated, and a highly potent FASN inhibitor in clinical trials for cancer (44) and non-alcoholic fatty liver disease (45). A commercially available related compound, TVB-3166, previously showed an IC50 in vitro of 42 nM and a cellular IC50 of 60 nM (46, 47).

Since we wanted to achieve complete inhibition of FASN, we used higher concentrations of TVB-3166. Treating cells with 0.2 μM or 20 μM TVB-3166 did not impact cell viability, nor did it alter the expression of Spike-C9 in HEK293T cells (Fig 6A, B). However, treating HEK293T cells expressing Spike-C9 with TVB-3166 or 2-bromopalmitate (2BP), a non-specific inhibitor of ZDHHC enzymes and other enzymes involved in fatty acid metabolism (48), attenuated the S-acylation of the Spike protein as determined by the APE assay (Fig. 6 C, D). Additionally, we examined whether TVB-3166 is a direct inhibitor of ZDHHC5 by assessing the covalent attachment of 15-HDYA on the Spike protein in the presence of TVB-3166. Consistent with Fig S3, we find that the exogenously added clickable fatty acid is incorporated more efficiently in the presence of the FASN inhibitor (Fig 6E, F), suggesting that there is competition between the endogenous acyl-CoAs and exogenous acyl chains. This also brings into question the mode of action of 2BP and whether it truly needs to be converted to 2BP-CoA to inhibit ZDHHC enzymes. Indeed, a previous study demonstrated that both 2BP and 2BP-CoA could directly modify Cys residues of ZDHHC enzymes and that 2BP can also irreversibly block Cys residues directly of target proteins such as Spike in our study (48). This further demonstrates the need for a better pan-inhibitor of ZDHHC enzymes.

**Figure 6.**
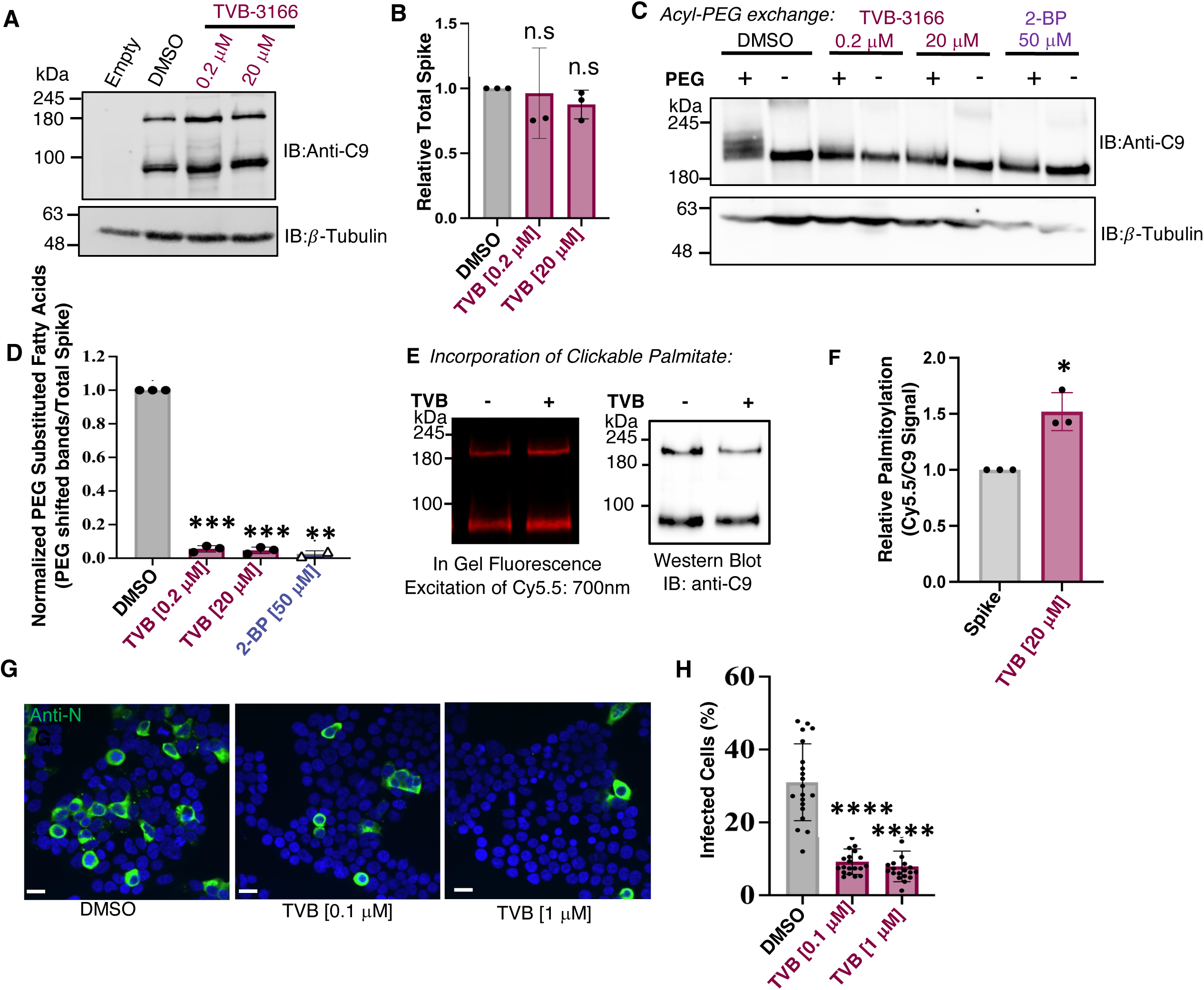
Inhibition of fatty acid synthase abolishes S-acylation of Spike and attenuates SARS-CoV2 spread *in vitro*. **(A)** Immunoblot of ectopically expressed Spike-C9 in HEK293T cells treated with DMSO, 0.2 μM or 20 μM TVB-3166 for 16-18 hrs. **(B)** Quantification of panel ‘A’. Data are the mean ± SEM n= 3 biological replicates normalized to GAPDH and analyzed using an unpaired t-test with Welch’s correction: 0.2 μM P=0.8637 and 20 μM P=0.1243. Inhibition of fatty acid synthase does not grossly impact the stability of the Spike protein. **(C)** Acyl-PEG exchange assay of Spike-C9 ectopically expressed in HEK293T treated with 0.2 μM or 20 μM TVB-3166 or 50 μM 2-Bromohexadecanoic acid (2-BP) for 16-18hrs. **(D)** Quantification of panel ‘C’ where PEG-modified bands were divided by the unmodified bands at ∼180 kDa, multiple unpaired t-tests with Welch’s correction, *** p=0.0001, ** p=0.0088 relative to DMSO control. Data are mean ± SEM of n= 3 biological replicates. **(E)** In-gel fluorescence (left) and immunoblotting (right) of Spike-C9 incubated with 15-HDYA following the treatment of DMSO or 20 μM TVB-3166 for 18h. Samples are processed as in Figure 1B. **(F)** Quantitation of panel ‘E’. Data are the mean ± SEM, n=3 replicates and analyzed using an unpaired t-test with Welch’s correction, * p=0.0334. **(G, H)** Representative images and quantification of HEK293T A2T2 cells (stably expressing Ace2 and TMPRSS2) infected with SARS-CoV2 (strain SB3, MOI of 0.1), treated with either 0.1 μM or 1 μM TVB-3166 6hrs post-infection for 18 hours. Data are the mean ± SEM from n = 3 replicates with all microscope fields plotted as individual points. Data were analyzed using a one-way ANOVA, **** P<0.0001 relative to DMSO. Scale bar = 6 μm.

To evaluate the effect of TVB-3166 on the viral spread, HEK293 A2T2 cells were infected with SARS-CoV-2 strain SB3 (49) at a MOI=0.1 and then treated with TVB-3166 6 hrs post-infection for an additional 18 hours. Then, cells were fixed, permeabilized, stained for the SARS-CoV-2 nucleocapsid protein, and imaged with confocal microscopy. Notably, a ∼70% decrease in infection was observed with TVB-3166 treatment (Fig 6 G). These experiments collectively suggest that inhibiting *de novo* lipid synthesis is an efficient way of attenuating S-acylation and that this strategy effectively blocks the in vitro spread of the SARS-CoV2.

### TVB-3166 mediated inhibition of FASN attenuates human CoV-229E spread and extends the survival of mice infected with a murine CoV

We confirmed that TVB-3166 treatment also inhibited the S-acylation of the 229E Spike-C9 protein. HEK293T cells were transiently transfected with 229E Spike-C9 and treated with TVB-3166; cell lysates were then subject to the ABE assay and revealed a decrease in the higher molecular weight species, thus poly-acylation (Fig. 7A, B). We next measured the ability of 229E to form plaques following TVB-3166 treatment. As illustrated in Fig 7 C and D, treatment with 0.2 μM TVB-3166 attenuated the spread of 229E by ≈ 85%. Together, these results suggest that the TVB-3166 FASN inhibitor block CoV infection at least in part by attenuating Spike S-acylation.

**Figure 7.**
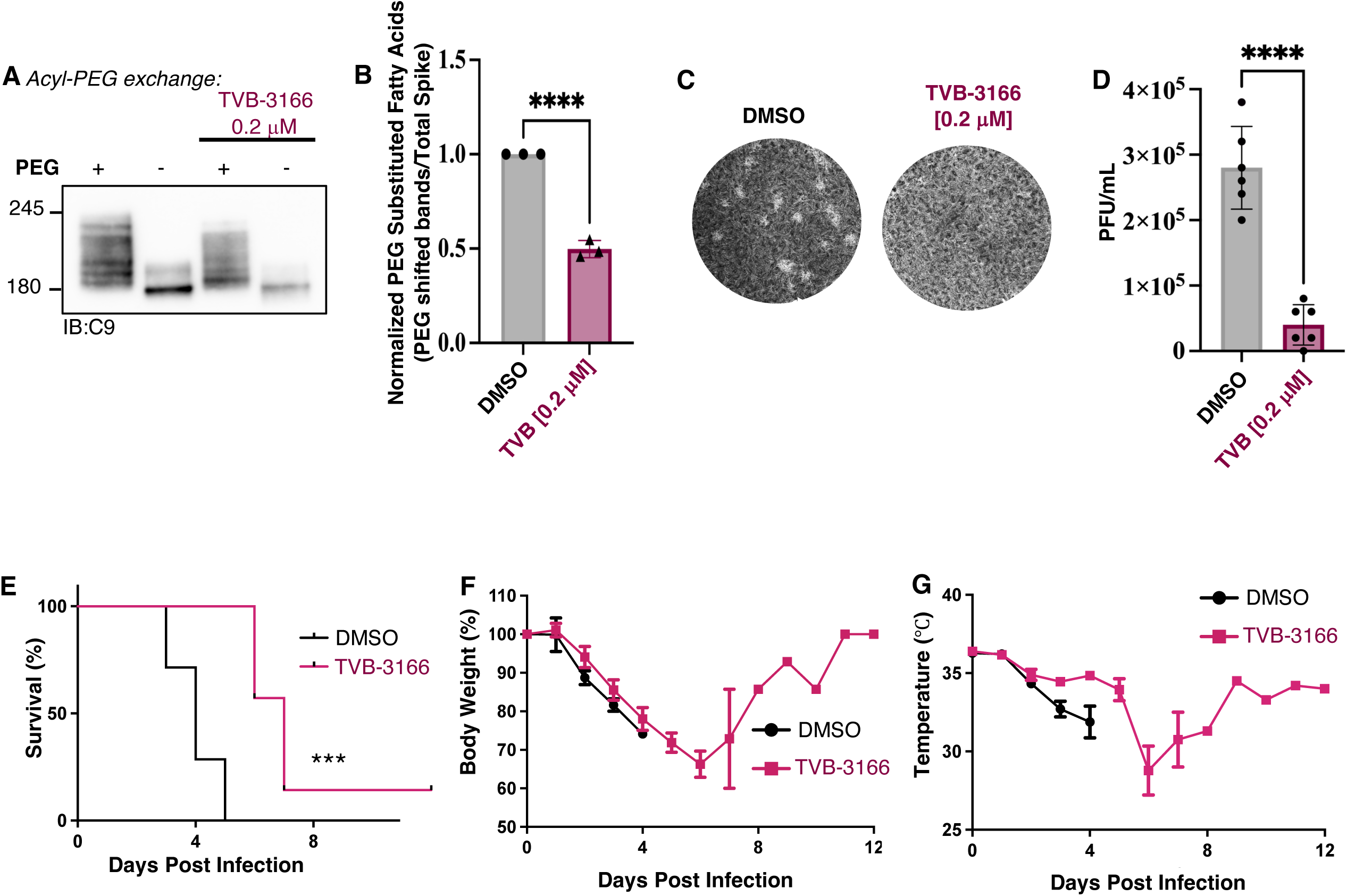
FASN inhibitor TVB-3166 attenuates the virulence of human and murine coronaviruses *in vitro* and *in vivo.* **(A)** Acyl-PEG exchange assay and quantification of 229E Spike ectopically expressed in HEK293T treated with 0.2 μM TVB-3166 for 16-18hrs. **(B)** Analysis of acyl-PEG assay where the PEG-modified bands normalized to unmodified bands at ∼180 kDa, unpaired t-test with Welch’s correction; data are mean ± SEM of n=3 biological replicates, **** p<0.0001. **(C, D)** Plaque assay of MRC-5 cells infected with 229E and treated with 0.2 μM TVB-3166 for 4 days, data are mean ± SEM of n=3 biological replicates, unpaired t-test**** P<0.0001. **(E)** Survival of mice challenged by a lethal dose of MHV-S and treated with TVB-3166 (30 mg/kg) following infection (DMSO group: n=7; TVB-3166 group n=7), survival curves analyzed using a Gehan-Breslow-Wilcoxon test. p < 0.005. **(F)** Bodyweight of mice challenged by a lethal dose of MHV-S and treated with TVB-3166 (30 mg/kg) following infection (DMSO group: n=7; TVB-3166 group n=7). DMSO treated mice were sacrificed around 4 d.p.i as demonstrated in ‘E’. **(G)** The temperature of mice challenged by a lethal dose of MHV-S and treated with TVB-3166 (30 mg/kg) following infection (DMSO group: n=7; TVB-3166 group n=7). DMSO treated mice were sacrificed around 4 d.p.i as demonstrated in ‘E’.

The cellular results demonstrating that TVB-3166 attenuated Spike S-acylation and limited spread of the SARS-CoV2 and 229E virus in cell-based assays motivated us to determine if this orally available FASN inhibitor could be useful as prophylactic in a mouse model. For this juvenile A/J mice were thus infected intranasally with 150,000 tissue culture infectious dose 50% of murine hepatitis virus (MHV)-S. In control mice, changes in health were noticed as early as two days post-infection; all proceeded to deteriorate as monitored by several health indicators, including decreased temperature, body weight, and activity (Fig 7 E-G). By day 5, all control mice were sacrificed as per animal care protocols. Previous experiments demonstrated that TVB-3166 is well tolerated by mice up to 100 mg/kg. As these are juvenile mice and the drug was delivered by oral gavage, we used a significantly lower dose of 30 mg/kg so as to not deliver too much fluid. Additionally, to help mitigate any putative metabolic issues, we also delivered the drug with a bolus of corn oil. In contrast to untreated mice, the animals treated daily with 30 mg/kg of TVB-3166 showed significantly prolonged survival (Fig 7E), with one animal recovering fully in this study. Thus, using a low dose of TVB-3166 at the time of infection, similar to a prophylaxis drug regimen, limited the progression of disease caused by MHV-S *in vivo* as marked by reduced clinical symptoms in treated mice. However, further studies are required to understand if the TVB compounds undergoing Phase II clinical for other diseases (50, 51) would be a beneficial treatment for CoV infections.

## Discussion

We and others have now established that the Spike protein of SARS-CoV2 is S-acylated. This modification supports its ability to catalyze membrane fusion and thus the infectivity of the SARS-CoV2 virus and experimentally useful pseudovirus (15,18,20). Furthermore, these observations are similar to previous findings on MHV and SARS Spike proteins (21, 23), demonstrating the importance of post-translational lipidation for the function of these fusogenic proteins. Collectively, various studies have now found that nearly half of the ZDHHC enzymes have the potential to S-acylate the Spike protein. Yet, there are no ZDHHC inhibitors, broad spectrum or selective, in clinical development. However, the findings that FASN inhibitors effectively block S-acylation of SARS-CoV2 Spike protein, attenuating the spread of the virus and even inhibiting the spread of other respiratory viruses, including respiratory syncytial virus, human parainfluenza 3 and rhinovirus (52) make this an area ripe for future investigations.

Mechanistically, the techniques that require cleavage of the thioester bond to detect sites of S-acylation, including acyl-biotin exchange, acyl-PEG exchange, and acyl-RAC, do not provide information as to the acyl chain species attached to the Cys residue(s). Using three alkyne-containing fatty acid analogs, we find that analogs of palmitate, myristate and stearate are attached to the Spike protein with a preference for the 16-carbon analog. We also found that ZDHHC5 has a strong preference for palmitoyl-CoA as a substrate and that when expressed in *ZDHHC5* knockouts, the incorporation of palmitate was reduced by 40%. The recent studies on the S-acylation of the Spike protein have relied on a variety of cell lines and techniques to examine post-translational lipidation. Therefore, the choice of cell types and different techniques may be contributing to the heterogeneity of the collective results. Alternatively, with its ten Cys residues, it’s also conceivable that the Spike protein is a promiscuous substrate for ZDHHC enzymes.

Considering the Spike protein contains ten Cys residues within the C-terminal domain and proximal to the transmembrane domain, it is worth considering the critical modification sites. A consensus from the various studies is that Cys1235 and 1236 are crucial for the infectivity of both the virus and pseudovirus. Additionally, a trend from these studies is that Cys1235, 1236, 1240 and 1241 are sites most highly modified by exogenously added ^3^H-palmitate or the clickable stearate analog 17-ODYA. Additionally, the results suggest that Cys1235 and 1236 are S-acylated in the endoplasmic reticulum by the ZDHHC20 before transiting to the Golgi apparatus and plasma membrane, where it can encounter additional ZDHHC isoforms. However, these studies rely on the uptake of fatty acids from the medium. The growing body of literature has demonstrated FASN, and not exogenous fatty acids, is the primary source of acyl chains for S-acylation (40,46,53–56). Using an alternative approach, Zeng et al. replaced all ten Cys with Alanine residues and subsequently re-introduced individual Cys residues within the tail and examined the S-acylation using the acyl-biotin exchange. This study found that the individual Cys1235, 1236, 1240 and 1241 residues were not being S-acylated in isolation, yet the other six sites were being modified (18). The significance of this finding is currently unclear. Still, it suggests that the recognition of the sites of S-acylation by the ZDHHC is more complex than just their proximity to the transmembrane domain. Another possibility is that thioesterases may more readily recognize some sites, especially in the absence of the other neighbouring S-acylated residue, so their steady-state S-acylation may appear lower. Indeed, how a de-acylation/re-acylation cycle may impact the findings of all these studies should be considered in the future.

This study, our recent studies on the peptidoglycan sensors NOD1 and NOD2 (40), and various other studies have demonstrated that the inhibition of FASN greatly attenuates the S-acylation of proteins even in the presence of exogenous lipids and lipoproteins (53,56,57). This observation is somewhat unexpected, although, to our knowledge, the source of fatty acids/fatty acyl-CoAs has not been rigorously examined. Treating cells with triacsin C a molecule known to block fatty acid uptake, does limit the uptake and incorporation of radiolabeled palmitate or clickable fatty acids from the medium (58, 59) as well as prevent the formation of lipid droplets. However, these studies did not investigate whether the proteins were still S-acylated by an independent method like the acyl-biotin exchange. Structurally unrelated FASN inhibitors, including the TVB compounds and cerulenin/C75, have comparable results; thus, this is unlikely the result of off-target effects. On the contrary, as we show in Fig 6F Fig S3, in the presence of TVB-3166, we see a tendency towards greater incorporation of the exogenously added clickable ligand.

One possible explanation is that exogenously added fatty acids are not internalized and converted to acyl-CoAs efficiently enough to supply the needs of the ZDHHC enzymes. Another possibility is that different acyl-CoA “pools” exist within the cytosol and that these may be channeled for further use and metabolism. Considering acyl-CoAs support phospholipid metabolism, triglyceride synthesis, protein acylation and can be imported into mitochondria and peroxisomes for β-oxidation, it may be difficult to resolve this question. Additionally, considering the concentration of CoA in the mitochondria is nearly two orders of magnitude higher than the cytosol(60), directly analyzing cytosolic CoA pools is not trivial. Our findings that the Spike protein can also be myristoylated and stearoylated (Fig. 4F), yet FASN inhibition abolished S-acylation using the APE assay (Fig. 6C), suggests that the FASN is also critical to generating the CoA species used for these reactions. In support of this notion, two recent studies have found that FASN generates an array of fatty acyl chains than just 16:0 and that FASN supports the generation of myristoyl-CoA and stearoyl-CoA and their subsequent attachment to proteins (61, 62).

Does the loss of ZDHHC5 and FASN activity impact the SARS-CoV2 infection cycle by other mechanisms? In all likelihood, yes. First, S-acylation may be important for the function of additional viral proteins. For comparison, the Envelope protein of MHV also requires S-acylation to assemble virions (63). Although we did not pick up other SARS-CoV-2 with the Acyl-RAC mass spectrometry analysis, it does not rule out the possibility that other viral proteins are, in fact, S-acylated. Indeed, in a previous study, we found it difficult to detect many of the SARS-CoV2 proteins, including those that could be potentially S-acylated (64). Second, loss of ZDHHC5 will alter S-acylated proteins at the cell surface, including flotillin, and is also known to increase the rates of two actin-dependent endocytic processes, phagocytosis and macropinocytosis, through an unknown mechanism (38, 40). Considering SARS-CoV2 is larger than the traditional clathrin-mediated endocytic cargo, other factors such as flotillin or actin machinery may play a role and thus be regulated by ZDHHC5. Additionally, since many proteins involved in vesicular transport and membrane fusion are S-acylated (65), including SNAP23 and SNARE proteins, treatment with FASN inhibitors would also impede these processes and thus attenuate the viral replication cycle. Indeed, many small molecules that interfere with phosphoinositides in the endocytic pathway and lysosomal pH are also known to disrupt the viral replication cycle (66–68).

Alterations in lipid metabolism are documented in CoV infections. Sera from patients with COVID19 have altered apolipoproteins and lipid levels (69), while 229E infected cells have been shown to have increased levels of free fatty acids (69). Given the fact that CoVs are enveloped viruses, they will require host lipids for replication. Indeed, CoV-induced remodeling of the cellular lipidome is necessary for robust viral replication. Specifically, recent studies have demonstrated that FASN activity is required to support SARS-CoV2 replication and that blocking fatty acid absorption using Orlistat reduced illness and symptoms in mice (67, 70). Additional experiments have demonstrated that the diacylglycerol acyltransferase 1 inhibitor (A922500) attenuates the production of viral progeny (71). This enzyme is critical for lipid droplet formation, and the same study found that SARS-CoV2 replications centers are in proximity to lipid droplets. Finally, the ectodomain of SARS-CoV2 Spike contains a binding pocket for another fatty acid, linoleic acid (72), although the function is not completely clear. Thus, further studies on the importance of fatty acids and lipid metabolism are needed to clarify these aspects of CoV infections.

Treating cells or mice with TVB-3166 limited the ability of SARS-CoV-2 and 229E to produce virulent progeny and spread in cell culture and in initial experiments extended the life of mice subjected to a lethal MHV infection. We anticipate that our findings will advance the possibility of using the related TVB-2640 that is currently undergoing clinical trials to treat COVID-19 or other CoV infections, especially as new variants arise (45, 73). Since FASN is a host enzyme, the likelihood of viral resistance is low. It would necessitate extensive mutagenesis of the Spike protein to bypass the requirement for S-acylation of the cytosolic tail to support membrane fusion. Clinically, the identification and approval of an orally available pan-CoV treatment or prophylaxis would be beneficial in the short term and against other zoonotic CoVs in the future.

## Acknowledgments

The authors would like to thank the Kennan Research Center for Biomedical Science Core Facilities at St. Michael’s Hospital, especially Dr. Caterina Di Ciano-Oliveira and Dr. Monika Lodyga, for their continued support, technical advice, expertise and training.

## Data and materials availability

All data is available in the main text or supplementary materials.

## Experimental procedures

### Reagents

TVB-3166 (SML1694), 2-Bromohexadecanoic acid (238422), EDTA-free protease inhibitor (1183617001), TCEP (C4706), tert-butanol (471712) were purchased from Millipore-Sigma (Oakville, Canada). Phusion polymerase, XhoI and NheI FastDigest restriction enzymes, T4 DNA ligase, dithiobissuccinimidyl propionate (DSP), Protein G Sepharose, Lipofectamine 3000, RNAiMAX, Eagle’s Minimum Essential Medium (EMEM), SuperSignal West Pico were purchased from Thermo Fisher Scientific (Mississauga, Canada). Dulbecco’s modified Eagle’s medium (DMEM), Temin’s Modified Eagle Medium (11935046), fetal bovine serum (FBS), charcoal stripped FBS, and Trypsin/EDTA were obtained from Wisent Bio Products (St. Bruno, Canada). Triton-X100, DMSO and CuSO_4_ were from BioShop Canada Inc (Burlington, Canada), alkynyl-palmitate (Click Chemistry Tools; Scottsdale, AZ), Cy5.5 (tetramethylindo)(di)- carbocyanines; Cyanine)-azide (Lumiprobe; Hunt Valley, Maryland), Site Counter (Badrilla; Leeds, UK), PFA (Electron Microscope Science; Hatfield, PA),

### Plasmids

pcDNA3.1-SARS2-Spike (Addgene plasmid #145032), pCEP4-myc-ACE2 (Addgene plasmid #141185). PBcmv-229E-wt_WRPE(39), HA-tagged ZDHHC5(40), EGFP-C1 (Clontech), mCherry-C1 (Clontech). The 229E Spike cDNA was amplified by PCR with the forward primer: 5’-TTTTTTGCGGCTAGCATGTTCGTGCTGCTGG and the reverse primer: 5’-TTTTTTGCGCTCGAGTCACGCCGGCGCCACCTGGCTGGTTTCGGTCTGGATGTGGATCTTTTCCA using Phusion polymerase (New England Biolabs) according to manufacturer’s instruction. The amplified 229E sequence and SARS-CoV2 S plasmid were incubated with XhoI and NheI restriction enzymes at 37°C for 1 hour. The reactions were then heat-inactivated and purified with a PCR clean-up kit. The insert and vector were then incubated with T4 DNA ligase at 16°C overnight. The ligation mixture was transformed into NEB 5-alpha high-efficiency *E. Coli* and plated on LB plates with the appropriate antibiotic. Plasmids were then harvested with a midi prep kit following the manufacturer’s instructions.

### Mutagenesis of SARS CoV-2 Spike multi-cysteine to serine

The SARS-CoV-2 Spike protein has ten cysteines in its cytosolic tail; C1235, C1236, C1240, C1241, C1243, C1247, C1248, C1250, C1253, C1254. In this study, all 10 Cys residues were replaced by Ser. Sequential PCR and overlap extension PCR was used for the complete substitution strategy. Using pcDNA3.1 SARS CoV-2 Spike-C9 as an initial template, two fragments were generated: 5’-and 3’-terminal fragments. To amplify the 3’-terminal fragment (containing 10 Cys to Ser mutation), three sequential PCR reactions were performed using overlapping forward primers (F2 primer overlaps with F1 primers, F3 primer overlaps with F2 primers). As a result, fragment 1 served as a DNA template for fragment 2, and fragment two as a template for fragment 3. The fragments 1,2 and 3 were generated using forward primersF1-5’-CTCCTCCTCCTCCGGCAGCTCCTCCAAGTTCGATGAGGACGATAG-3’, F2-5’-CCTCCTCCTCCAGCTCCCTGAAGGGCTCCTCCTCCTCCGGCAGCT-3’, and F3-5-TGATGGTGACCATCATGCTGTCCTCCATGACCTCCTCCTCCAGCTCCCTG-3’, respectively, and reverse primer 5’-TCTAGACTCGAGCTAAGCGGGAGC-3’. To amplify the 5’ DNA fragment, forward primer 5’-CAAGCTGGCTAGCATGTTTGTCTTCC-3’ and reverse primer 5’-TCATGGAGGACAGCATGATGGTCACCATCA-3’ were used. The 5’ DNA fragment and 3’DNA fragment were annealed together with their complementary overhanging by PCR using the primer pairs 5’-CAAGCTGGCTAGCATGTTTGTCTTCC-3’ and 5’-TCTAGACTCGAGCTAAGCGGGAGC-3’ to generate full-length DNA. All PCR thermocycler conditions were based on Touchdown PCR(74). The multi-site mutated SARS CoV-2 Spike was then subcloned into the pcDNA3.1 vector using NheI and XhoI restriction digestion.

### Cell culture

HEK293T cells were cultured in DMEM supplemented with 10% FBS at 37°C and 5 CO_2_. MRC-5 cells (ATCC) were cultured in EMEM supplemented with 10% FBS and 1% Penicillin/Streptomycin (Gibco).

### Syncytium formation assay

HEK293T cells were seeded in 6-well plates and transfected with 1. EGFP-C1 vector with myc-ACE2 vector and 2. mCherry-C1 vector with CoV-2 Spike-C9 or Spike multi-cysteine to serine mutant. After 16-24 hrs of incubation, cells were lifted by trypsinization and co-cultured overnight. Fluorescent images were acquired through EVOS FLoid™ Cell Imaging System.

### Antibodies and antibody-conjugated beads

C9 antibody (Clone 1D4, Santa Cruz, sc-57432), anti-C9 agarose bead (Cube Biotech), HA antibody (Abcam, ab9110), SARS-CoV-2 (COVID-19) Spike RBD antibody (GenTex, HL257), ZDHHC5 antibody (Sigma, HPA014670), fluorescent secondary antibodies (Jackson Laboratories, Invitrogen, LI-COR).

### siRNA Transfection

MRC-5 cells seeded on 6-well tissue culture plates were transfected 2X, 24 hours apart, followed by a 24 to 48-hour recovery before infection with 229E. Cells were initially transfected at 50% confluence in 1mL Opti-MEM Reduced-Serum Medium (Gibco). A siRNA transfection master mix of Lipofectamine RNAiMAX, siRNA (CTRL or ZDHHC5), and Opti-MEM was made according to the manufacturer’s instructions, with a final siRNA concentration of 50nM. Cells were transfected with the siRNA master mix for 4 hours, followed by a change to growth media. After the last transfection, media was changed to growth media and cells were given 24 to 48-hour recovery before the infection, at which point they were at 100% confluence. Oligo sequences used CTRL non-targeting siRNA: CGUACUGCUUGCGAUACGGUU and ZDHHC5 siRNA: CUGUGAAGAUCAUGGAUAAUU (38).

### Immunoblotting

SDS polyacrylamide gels were transferred on PVDF membranes by Trans-blot Turbo System at 25 Volts for 20 min. Membranes were blocked with 5% BSA for 1 hour at room temperature and were incubated with primary antibodies diluted in blocking solution at 4°C overnight (anti-C9 Santa Cruz). Membranes were washed with TBS-0.1% Tween20 (TBST) for 5 minutes, three times, and incubated with appropriate secondary antibodies. Blots were then washed three times with TBST for 5 minutes each wash. The membranes were treated with ECL and imaged with ChemiDoc Imaging System (BioRad).

### Co-Immunoprecipitation

HEK293T cells seeded in T25 Flask (25cm^2^) were grown until reaching 70-90% confluency. According to the manufacturer’s instruction, cells were transfected with 3xHA-zDHHC5 and CoV-2 Spike-C9 plasmids using Lipofectamine 3000 and incubated for 16-24hrs. For protein cross-linking, cells were incubated with a 2mM DSP crosslinker in PBS at four °C for 2hrs. DSP was quenched in 20 mM glycine in PBS for 15 min, and cells were lysed in NP40 lysis buffer (20mM Tris and adjusted to pH 7.4, 150 mM NaCl, 1% NP40, two mM EDTA) supplemented with cOmplete Protease Inhibitor Cocktail tablet, EDTA-Free at a constant agitation for 30 min at 4°C. Cell lysates were centrifuged at 12,000xg for 20 min, and the supernatant was collected. Samples were pre-cleared by incubating with protein G Sepharose beads and were spun at 14,000 xg for 1min to collect the supernatant. Pre-cleared samples were incubated with anti-C9 or anti-HA primary antibodies overnight at 4°C. To form a primary antibody-bead complex, Protein G beads were added and incubated for 30min. Beads were washed in cold wash buffer (20mM Tris, pH 7.4, 350 mM NaCl, 1% NP-40, 2 mM EDTA) 6 times, and proteins were eluted with SDS Laemmli buffer and heated for 30min at 37°C before SDS-PAGE.

### Immunofluorescence

Cells were seeded on an 18mm-round coverslip in a 12-well plate. Cells expressing Spike-C9 or Spike^C->S^-C9 ectopically were washed with PBS and then fixed with 4% PFA. The remaining PFA was quenched by 0.15M glycine for 20 min at RT. Cells were blocked with 5% BSA in PBS for 1hr and probed with a primary antibody (SARS-CoV-2 Spike RBD, GenTex) in 5% BSA for 1hr. Cells were washed with PBS and probed with fluorescently labelled secondary antibodies for 1hr.

### Confocal Microscopy

Imaging in the St. Michael’s Hospital BioImaging was conducted using an Andor Diskcovery multi-modal imaging system provided by Quorum Technology (Guelph, Ontario). The system is based on a Leica DMi8 equipped with a 63x/1.47 NA oil objective; 405, 488, 561, and 637 nm laser lines; 450/50, 525/50, 600/50, 610/75, and 700/75 emission filters. Images were acquired using Metamorph software on a Hamamatsu ORCA-Flash 4.0 V2 sCMOS camera. Imaging at Ryerson University was performed on the Quorum WaveFX spinning disc based on an inverted Leica DMi8 equipped with an Andor Zyla 4.2 sCMOS camera using a 10x objective controlled by MetaMorph.

### Metabolic labelling/ on-bead click chemistry assay

HEK293T cells seeded in T25 Flask (25cm^2^) were transfected with pcDNA3.1-SARS CoV-2-Spike-C9 and incubated overnight in DMEM supplemented with 10% (v/v) charcoal-stripped FBS with or without 100 μM of clickable fatty acid analogs (Click Chemistry Tools, Scottsdale, AZ) coupled to albumin. After 18 hours of labelling, cells were washed twice with ice-cold PBS and collected following scraping. Cells were lysed in IP buffer (20 mM Tris and adjusted to pH 8.0, 137 mM NaCl, 1% NP40, 2 mM EDTA) supplemented with EDTA-free cOmplete Protease Inhibitor Cocktail tablet (Roche) and immunoprecipitated using C9-agarose bead (Cube biotech) at 4°C overnight. The beads were then washed with ice-cold D-PBS 4 times and incubated in D-PBS supplemented with 1 μM Cy5.5-azide (Click Chemistry Tools), 1 mM TCEP (adjusted to pH 7.0), 1 mM CuSO_4_, 0.1 mM TBTA (1:4 v/v DMSO: tert-Butanol) for 2h. The beads were washed 5 times with D-PBS, and proteins were eluted with 5X SDS Laemmli buffer. Protein samples were resolved by SDS-PAGE and subjected to in-gel fluorescence using the 685-700 nm laser on the Li-Cor Odyssey Infrared Imaging System. Gels were later transferred to PVDF membranes for immunoblotting, as previously indicated.

### Flow Cytometry

HEK293T cells seeded in T25 Flask (25cm^2^) were transfected with pcDNA3.1 SARS-CoV2 Spike-C9 or pcDNA3.1 SARS-CoV2 Spike-C9^C->S^ and incubated overnight in DMEM supplemented with 10% (v/v) FBS. Cells were washed with PBS and then gently dethatched with 10mM EDTA (pH=7). Cells were collected by centrifugation at 1,500 rpm for 5 mins and were then resuspended in flow buffer (PBS supplemented with 2% (v/v) FBS). 1x10^6^ cells were then immunolabeled with primary antibody (SARS-CoV-2 Spike RBD, GenTex) inflow buffer for 30 mins at 4(. Cells were washed three times with flow buffer and then incubated with a secondary antibody. Cells were washed in flow buffer, and DAPI was added to cells right before acquisition with the DB LSRFortessa^TM^ X-20 Cell Analyzer. Analysis was completed on the FLowJo software (BD Biosciences).

### Acyl-PEG exchange assay

HEK293T cells were seeded in 6 well plates in DMEM +10% FBS and were transfected with lipofectamine 3000 reagent. For each well 2.5 mg of either: 229E Spike-C9 + HA-ZDHHC5, 229E Spike-C9+ EGFP-C1, SARS-CoV-2 Spike-C9 and HA-ZDHHC5 or SARS-CoV2 Spike-C9 + EGFP-C1 DNA mixtures were diluted in Opti-mem and mixed with Lipofectamine 3000 according to the manufacturer’s instructions. Upon transfection, cells were treated with 0.2 μM or 20 μM of TVB-3166 or 50 μM 2-BP. 16 -18 hours post-transfection, cells were lysed and processed with the SiteCounter^TM^ S-palmitoylated protein kit (Badrilla, Leeds, UK) following the manufacturer’s instructions. Samples were subjected to SDS-PAGE by resolving on a 3-8% Criterion^TM^ XT Tris-Acetate gel (Bio-Rad) for immunoblotting.

### Propagation of the Human 229E Coronavirus

Human coronavirus 229E (ATCC^®^ VR-740^™^) was purchased from American Type Culture Collection (ATCC), Manassas, VA. Stocks of the 229E virus were produced by propagation in MRC-5 cells. Briefly, a T75 flask of 90% confluent MRC-5 cells was infected with an MOI of 0.01 for 2 hours at 33°C in infection media for viral adsorption. Unbound virus was removed by washing cells 2X with infection media, and infected cells were returned to the 33°C incubator in 12mL infection media for 72 hours. The virus-containing supernatant was centrifuged at 1000 xg for 5 minutes to remove cellular debris, and single-use aliquots of the virus were frozen at -80°C.

### Human CoV 229E plaque-forming unit assay

PFU assays were used to determine viral titers and experiments using protocols adapted from previous publications field(75, 76). Briefly, MRC- 5 cells seeded on 6-well tissue culture and grown to 100% confluency in EMEM+ 10% FBS+ 1% PenStrep. Plates were infected with serially diluted virus in infection media (EMEM+2% FBS+ 1% PenStrep) for 1 hour at 33°C in a volume of 300uL with periodic agitation. After virus adsorption, the unbound virus was removed by washing with infection media followed by agarose overlay with 1X MEM Temin’s modification (Thermo Fisher, 11935046) +0.3% agarose. Cells were then returned to the 33°C incubator until they showed obvious signs of cytopathic effects (typically 4-7 days post-infection), upon which plates were fixed with 10% neutral-buffered formalin for 4 hours, followed by removal of the agarose plug and staining with 1% crystal violet for 15 minutes. Plaques were counted based on standard protocols for determining viral titer.

### Liquid viral infection assay

MRC-5 cells were seeded on glass coverslips and transfected. Cells were infected with 229E at an MOI of 0.005 for 2 hours in a total volume of 2mL per well. After virus adsorption, cells were washed 2X with infection media and then replaced with 2mL infection for incubation at 33°C. After 72 hours, cells were fixed with 4% PFA for 1 hour, followed by: 0.15% glycine, 0.1% Triton X-100, and 3% BSA for 15 minutes each. Cells were then incubated in a 1:50 dilution of mouse anti-229E Spike protein antibody by inverted drop for 1 hour at room temperature. After washing, cells were incubated in 1:1000 anti-mouse Alexa Fluor 488 secondary antibody (Jackson ImmunoResearch) and DAPI (1 mg/mL) for 1h at room temperature, followed by mounting on slides with DAKO mounting media. Images were acquired with the WaveFX system. The total fluorescent signal was quantified with ImageJ.

### SARS-CoV-2 infection and immunofluorescence

293A2T2 (HEK293 expressing ACE2 and TMPRSS2 ectopically) cells were grown in DMEM media supplemented with 10% heat-inactivated FBS. 20,000 cells were plated on coverslips in a 24-wells plate the day before infection and infected with SARS-CoV-2 strain SB3(49) at an MOI of 0.1 in DMEM for 1hr. Infectious media was changed with DMEM-FBS2%, and cells were incubated at 37C at 5% CO2. For TVB treatment, at 6h post-infection, the supernatant was replaced with fresh media DMEM-FBS2% containing either DMSO or the TVB compound at 0.1 or 1 μM final concentration. Cells were incubated for 18h at 37°C, with 5% CO2. After infection, cells were fixed with 4% paraformaldehyde (PFA) for 1hour at room temperature. After PFA fixation, cells were washed with PBS and permeabilized for 10 min with 0.1% Triton X-100. Cells were blocked in 2% bovine serum albumin (BSA) in PBS for 1hour before incubation with the primary antibody overnight at 4C. Primary anti-nucleocapsid antibody (SARS-CoV-2 Nucleocapsid Antibody (HC2003), Cat. No. A02039, GenScript) was used at 1:1000. After removing the antibody solution, cells were washed in PBS and incubated for 1hour with secondary anti-human FITC (Biolegend) at room temperature. Cells were washed in PBS, and coverslips were mounted with Duolink *in situ* mounting medium with DAPI (Sigma, cat# DUO82040). Cells were imaged using UPlan FL 40x/0.75 NA on a Yokogawa CSU-10 Spinning-disk confocal microscope. Images were processed using the Volocity Viewer v.6.

### MHV-S infection of mice

Four-week-old female A/J mice were purchased from Jackson Laboratories. Mice were housed in the St. Michael’s Hospital Vivarium on a standard light: dark cycle and were given free access to food and water. Before infection, mice were acclimatized for one week. Mouse infection experiments were performed in the dedicated BSL-2 room in the vivarium in accordance with the St. Michael’s Hospital Animal Care Committee guidelines for animal use; approved animal protocol (ACC962, St. Michael’s Hospital). One day prior to infection, the necks of mice were shaved, and chemical depilatory cream was used to remove all hair to obtain optimal oxygen saturation measurements during the study. For infection, mice were sedated with 5% isoflurane and infected intranasally with 150,000 units determined using a tissue culture infectious dose at 50% (TCID_50_)/mouse of MHV-S (Cedarlane, VR-766), which is known to induce pneumonitis (77). The virus was diluted in PBS to a final volume of 75 μL before administration. After infection, mice were allowed to recover for ten minutes in a cage on a covered heating pad. On the day of infection, TVB-3166 (Sigma, SML 1694) was reconstituted by adding a small volume of DMSO (10% of the final volume) to the vial. The vial was briefly vortexed and heated in a 37°C water bath, and then corn oil was added to make a 5mg/mL solution for treatments. Following recovery, mice were randomly separated into weight-matched groups and immediately received solvent control or TVB-3166 (30 mg/kg) by oral gavage. For the remainder of the study, mice were monitored daily and received one dose of TVB-3166 or solvent control per day. As previously reported (78), mice were sacrificed if they reached two of four endpoints: weight loss of 30% of the initial weight, body temperature below 31°C, oxygen saturation below 75% or activity level of 1 (moribund). To measure oxygen saturation on awake mice, the Mouse Ox Plus device and software from Starr Life Sciences were used.

### Acyl resin-assisted capture (Acyl-RAC) assay

HEK293 A2T2 and African green monkey VeroE6 cells were infected with SB3 SARS-CoV-2 for 24 hours at an MOI of 0.5. Cells were harvested and lysed (1% Triton X-100, 50 mM Tris-HCl (pH 7.5), 150 mM NaCl, 1 mM EDTA, 1 % SDS, 0.5 % sodium deoxycholate, 100 µL mammalian protease inhibitor cocktail (Sigma #P8340), 5 µL Turbonuclease (BioVision # 9207-50KU), 10 mM N-ethylmaleimide (NEM, BioShop #ETM222.5)). Lysates were sonicated and incubated at 4°C on an end-over-end rotator for 1 h. Additional NEM was added to a final concentration of 20 mM and samples were incubated for an additional hour on an end-over-end rotator at room temperature. Next, NEM was removed via methanol chloroform precipitation, and protein pellets were rinsed with methanol and air-dried. Pellets were resuspended in Acyl-RAC binding buffer (100 mM Tris-HCl (pH 7.5), 1 mM EDTA, 1 % SDS) by sonication. Hydroxylamine (NH_2_OH, 400 mM final concentration; or NaCl for control samples) was added before incubation with 30 µL of packed acyl-RAC beads (High-Capacity Acyl-RAC S3 beads, Nanocs # AR-S3-1,2) for 2 hours at room temperature. Next, beads were washed 3 times with binding buffer and 10 times with NH_4_HCO_3_ (50 mM, pH 8.3). Direct digestion on beads was performed using trypsin/Lys-C (1 µg per sample in 50 mM NH_4_HCO_3_, pH 8.3) for 5 h at 37 °C on an end-over-end rotator. Beads were rinsed with 50 mM NH_4_HCO_3_ (pH 8.3), and supernatant fractions were pooled. Samples were desalted and lyophilized before LC-MS analysis.

### Liquid Chromatography-Mass Spectrometry

LC-MS analysis was performed as previously described (64). Briefly, HPLC was performed on samples reconstituted in HCOOH (0.1%) and loaded on a 20 mm pre-column (C18 Acclaim PepMap^TM^ 100, 75 µm x 2 cm, 3 µm, 100Å, Thermo Scientific) and separated on a 50 mm analytical column (C18 Acclaim PepMap^TM^ RSLC, 75 µm x 50 cm, 3mm, 100Å, Thermo Scientific) over a 2-hour reversed-phase gradient (5-30% CH3CN in 0.1% HCOOH) with 250 nl/min flow rate on an EASY-nLC1200 pump in-line with a Q-Exactive HF mass spectrometer (Thermo Scientific) operated in positive ion mode ESI. MS scans were performed at a resolution of 60,000 (fwhm), followed by up to 20 MS/MS scans (HCD, 15,000 fwhm) of the most intense parent ions. Dynamic exclusion (within 10 ppm) was set for 5 seconds.

### Mass Spectrometry Data Processing

For the sequence database search, Thermo raw files (.raw) were converted to the .mzML format using Proteowizard (v3.0.19311), then searched using X!Tandem (v2013.06.15.1) and Comet (2014.02 rev. 2) against Chlorocebus sabaeus RefSeq (GCF_015252025.1, v102; 61,745 entries) for the VeroE6 samples, or Human RefSeq (v104; 36,113 entries) for the HEK293-A2T2 samples. Both databases were supplemented with the SARS-CoV-2 proteome (2019-nCoC HKU-SZ-005b7). Search parameters specified a parent MS tolerance of 15 ppm and an MS/MS fragment ion tolerance of 0.4 Da, with up to two missed cleavages allowed for trypsin/Lys-C. Deamidation (NQ), oxidation (M), acetylation (protein N-term), N-ethylmaleimide (C), and palmitoylation (C) were set as variable modifications. Search results were processed using the trans-proteomic pipeline (TPP v4.7), and proteins with an iProphet probability ≥0.9 with at least two unique peptides matched were considered high confidence identifications. Bayesian statistics (SAINT)(79) were applied to compare NaCl-treated samples to hydroxylamine-treated samples in order to identify putative acylated proteins. Control (NaCl-treated) and hydroxylamine-treated replicates were compressed to 2 and a BFDR cut-off of 1% was used.

### Data availability

All MS data have been deposited on Mass Spectrometry Interactive Virtual Environment (massive.ucsd.edu) under accession MSV000088105. All other data is available upon request.

## Supporting information

This article contains supporting information. Three supplementary figures and two supplementary tables are included.

## Author contributions

Katrina Mekhail; methodology and formal analysis, writing. Minhyoung Lee; methodology and formal analysis, writing. Michael Sugiyama; Methodology and formal analysis, editing. Audrey Astori; methodology and formal analysis. Jonathan St-Germain; methodology and formal analysis. Elyse Latreille; methodology and formal analysis. Negar Khosraviani; formal analysis. Kuiru Wei; formal analysis. Zhijie Li; resources. James Rini; resources. Warren Lee; project administration, funding. Costin Antonescu; project administration, funding. Brian Raught; project administration, funding. Gregory Fairn; project administration, funding, writing, editing.

## Funding and additional information

This work was supported by the St. Michael’s Hospital Foundation and a Project Grant from the Canadian Institutes of Health Research (PJT166010) to GDF. WLL is supported by a Canada Research Chair in Mechanisms of Endothelial Permeability and operating funds from the Keenan Foundation, CHRP grant (CPG 158284; CHRP 523598) and a Grant from CIHR (FRN 170656). CA received funding from the Ryerson Faculty of Science and the Ryerson COVID-19 SRC Response Fund. ML is supported by a doctoral scholarship from the Natural Sciences and Engineering Research Council of Canada. EL is supported by a Canadian Graduate Scholarship – Master’s Program and a Scholarship from the St. Michael’s Hospital Research Training Center Scholarship.

## Supplemental Figures

**Supplemental Figure 1.**
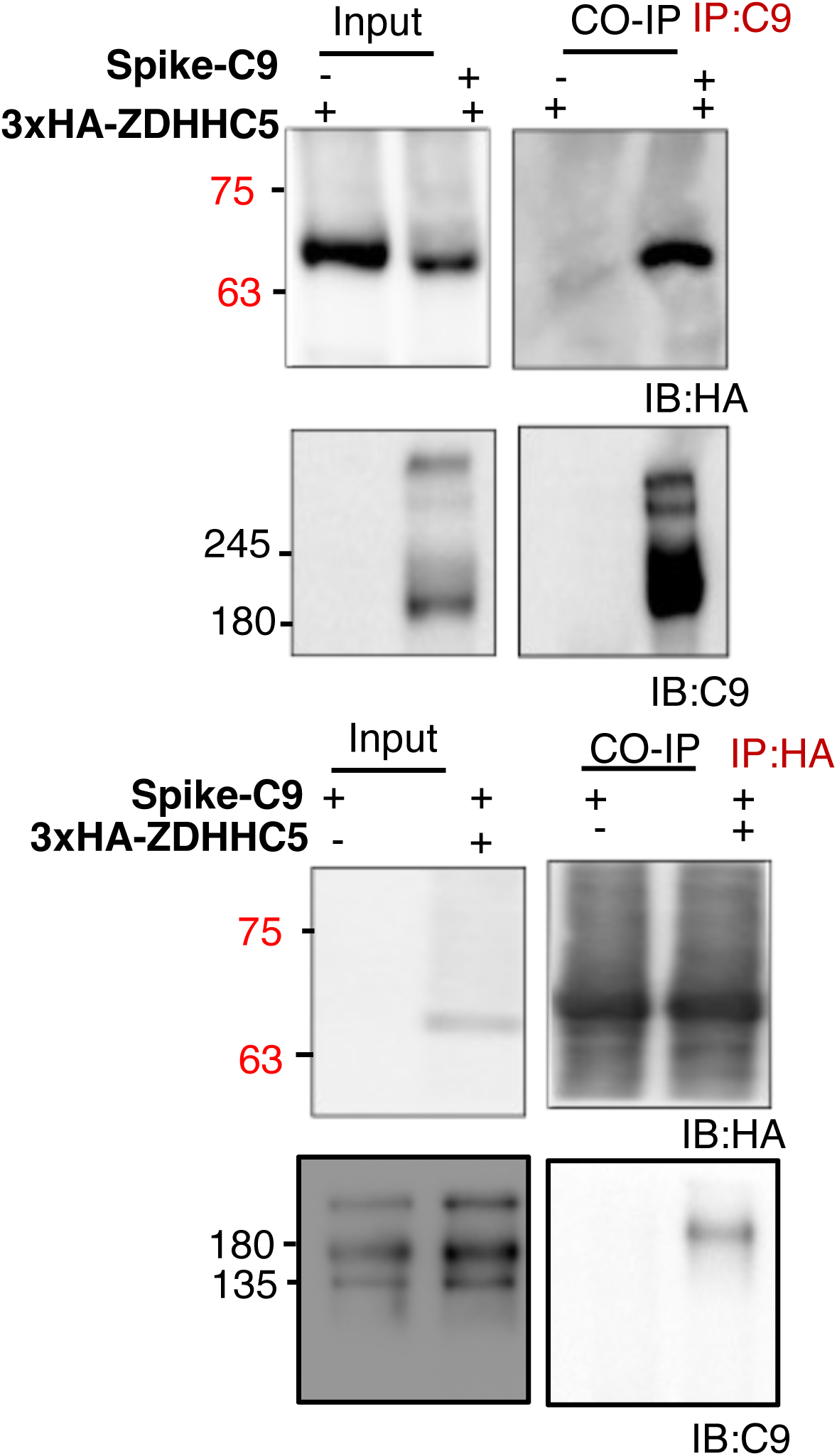
Co-immunoprecipitation of HA-ZDHHC5 and Spike-C9. Co-immunoprecipitation of SARS-CoV-2 Spike-C9 with 3xHA-ZDHHC5. HEK293T cells co-transfected with Spike-C9 and 3xHA-ZDHHC5 were immunocaptured and probed.

**Supplemental Figure 2.**
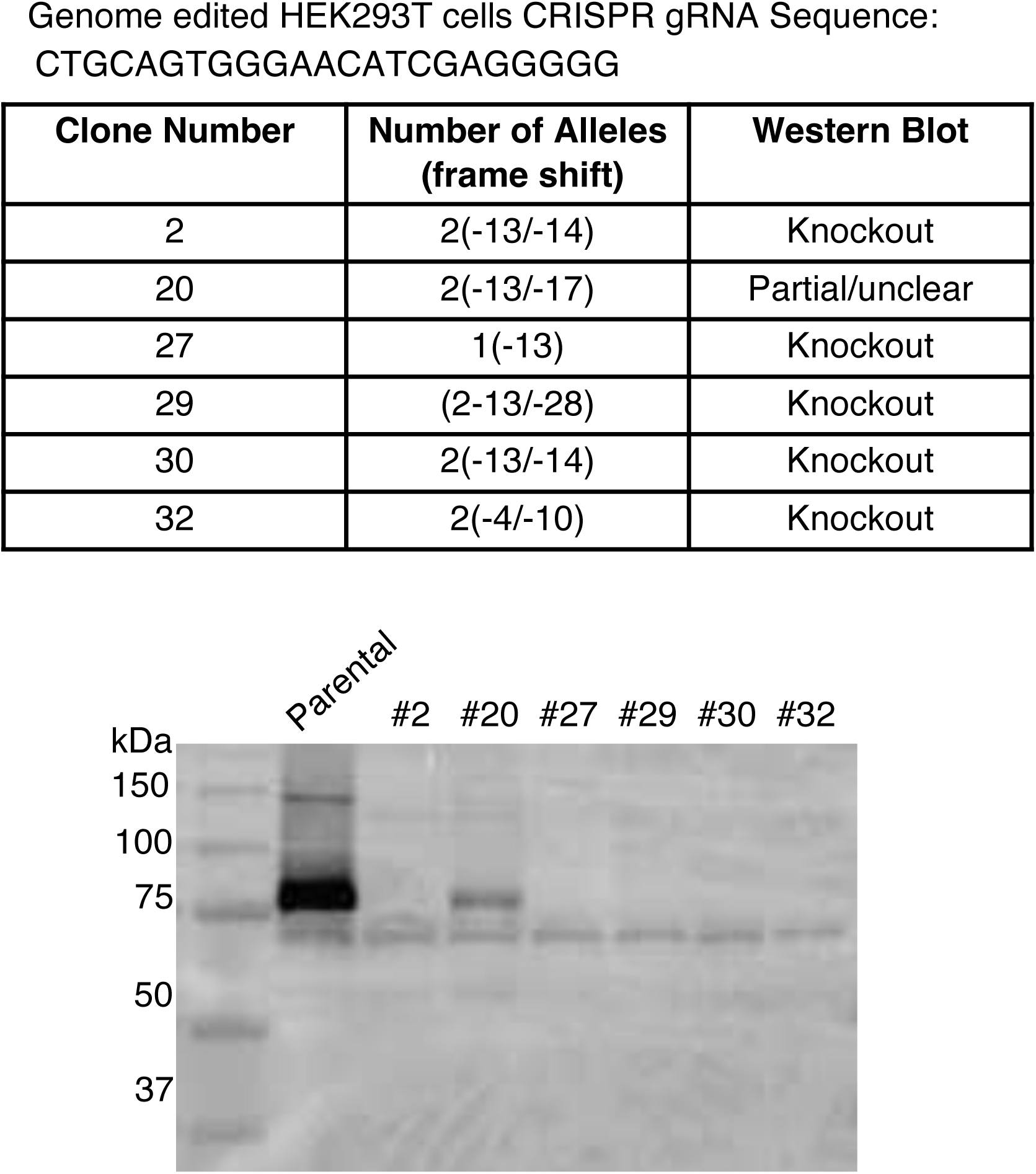
Generation of the ZDHHC5 knockout HEK293 Cells. Using the gRNA, nearly 50 clones were isolated and validated by sequencing to identify specific deletions and frameshifts. Clones containing deletions that result in frameshifts were subjected to immunoblotting.

**Supplemental Figure 3.**
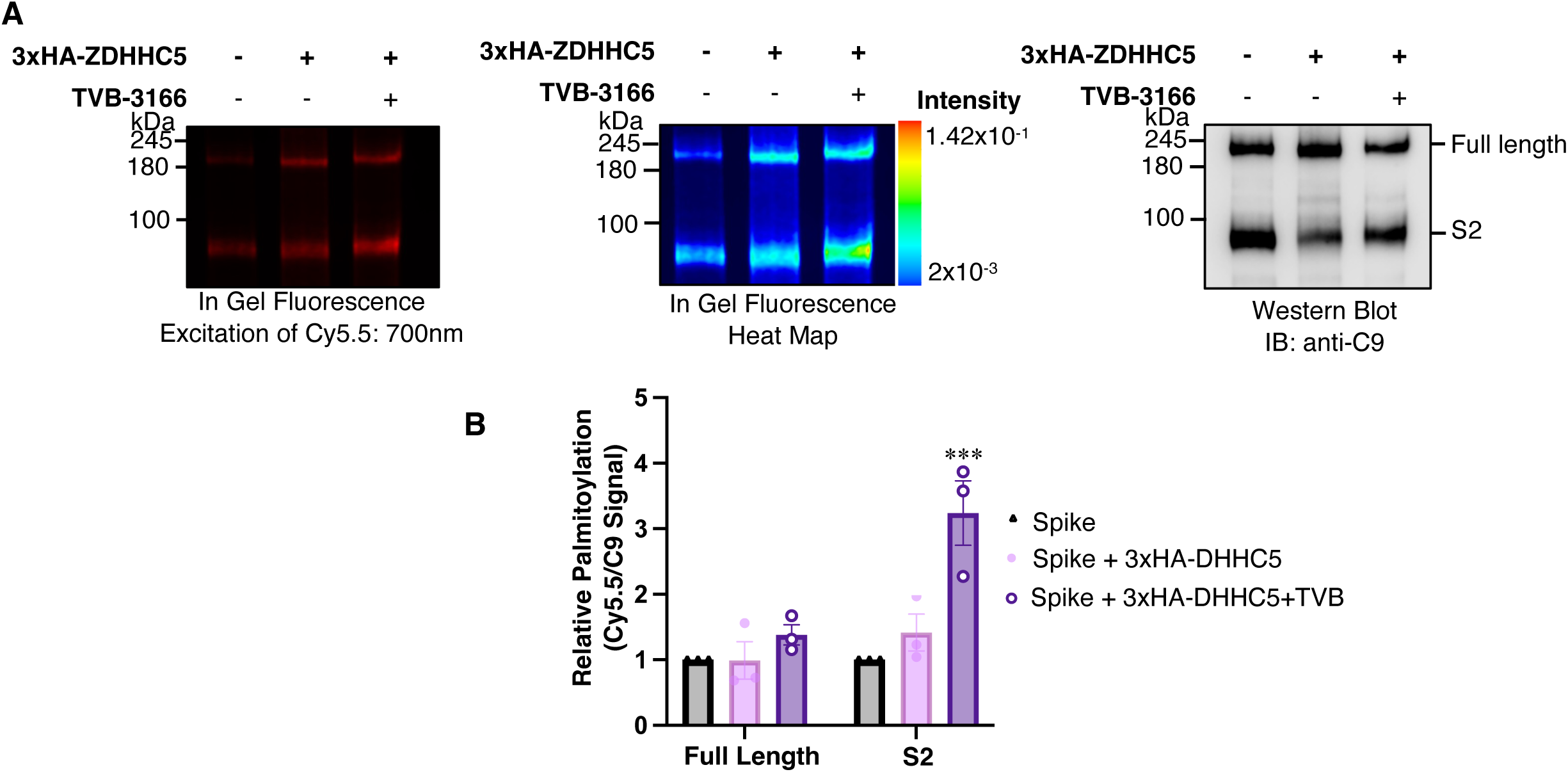
Inhibition of *de novo* synthesis increases the incorporation of clickable palmitate on Spike-C9. **(A)** In-gel fluorescence (left), a pseudo-coloured heat map (center) and anti-C9 immunoblot (right) of control HEK293T cells and cells transiently transfected with 3HA-ZDHHC5 with or without TVB-3166 for 18 hrs. Increased abundance of ZDHHC5 alone has a modest impact on the incorporation of 15-HDYA onto Spike-C9 while simultaneously inhibiting de novo fatty acid synthesis and increasing the attachment of the palmitate analog. **(B) Quantitation of panel ‘A’,** fold increase in Spike palmitoylation was determined by normalizing the amount of Cy5.5-Azide fluorescence to the amount of protein. Data represent the mean ± SEM, n = 3 biological replicates. Multiple unpaired t-tests, Spike vs. Spike + 3xHA-ZDHHC5, full length n.s P=0.977024, n.s P=0.214677, Spike vs Spike-C9+ ZDHHC5 +TVB3166; full length: n.s P=0.067377, S2 *** P=0.010351.

**Supplemental Table 1. Complete list of S-acylated proteins in Vero and HEK293T A2T2 cells**

**Supplemental Table 2. Peptide counts and protein info from the acyl-RAC mass spectrometry identification.**

## References

1. Wang, C., Horby, P. W., Hayden, F. G., and Gao, G. F. (2020) A novel coronavirus outbreak of global health concern. Lancet 395, 470–473

2. da Costa, V. G., Moreli, M. L., and Saivish, M. V. (2020) The emergence of SARS, MERS and novel SARS-2 coronaviruses in the 21st century. Arch Virol 165, 1517–1526

3. To, K. F., Tong, J. H., Chan, P. K., Au, F. W., Chim, S. S., Chan, K. C., Cheung, J. L., Liu, E. Y., Tse, G. M., Lo, A. W., Lo, Y. M., and Ng, H. K. (2004) Tissue and cellular tropism of the coronavirus associated with severe acute respiratory syndrome: an in-situ hybridization study of fatal cases. J Pathol 202, 157–163

4. Weiss, S. R., and Leibowitz, J. L. (2011) Coronavirus pathogenesis. Adv Virus Res 81, 85–164

5. Monteil, V., Kwon, H., Prado, P., Hagelkruys, A., Wimmer, R. A., Stahl, M., Leopoldi, A., Garreta, E., Hurtado Del Pozo, C., Prosper, F., Romero, J. P., Wirnsberger, G., Zhang, H., Slutsky, A. S., Conder, R., Montserrat, N., Mirazimi, A., and Penninger, J. M. (2020) Inhibition of SARS-CoV-2 Infections in Engineered Human Tissues Using Clinical-Grade Soluble Human ACE2. Cell 181, 905–913 e907

6. Hoffmann, M., Kleine-Weber, H., Schroeder, S., Kruger, N., Herrler, T., Erichsen, S., Schiergens, T. S., Herrler, G., Wu, N. H., Nitsche, A., Muller, M. A., Drosten, C., and Pohlmann, S. (2020) SARS-CoV-2 Cell Entry Depends on ACE2 and TMPRSS2 and Is Blocked by a Clinically Proven Protease Inhibitor. Cell 181, 271–280 e278

7. Kim, Y. C., Dema, B., and Reyes-Sandoval, A. (2020) COVID-19 vaccines: breaking record times to first-in-human trials. NPJ Vaccines 5, 34

8. Hoffmann, M., Kleine-Weber, H., and Pohlmann, S. (2020) A Multibasic Cleavage Site in the Spike Protein of SARS-CoV-2 Is Essential for Infection of Human Lung Cells. Mol Cell 78, 779–784 e775

9. Duffy, S., Shackelton, L. A., and Holmes, E. C. (2008) Rates of evolutionary change in viruses: patterns and determinants. Nat Rev Genet 9, 267–276

10. Duffy, S. (2018) Why are RNA virus mutation rates so damn high? PLoS Biol 16, e3000003

11. Cui, J., Li, F., and Shi, Z. L. (2019) Origin and evolution of pathogenic coronaviruses. Nat Rev Microbiol 17, 181–192

12. Liu, L., Fang, Q., Deng, F., Wang, H., Yi, C. E., Ba, L., Yu, W., Lin, R. D., Li, T., Hu, Z., Ho, D. D., Zhang, L., and Chen, Z. (2007) Natural mutations in the receptor binding domain of spike glycoprotein determine the reactivity of cross-neutralization between palm civet coronavirus and severe acute respiratory syndrome coronavirus. J Virol 81, 4694–4700

13. Gordon, D. E., Jang, G. M., Bouhaddou, M., Xu, J., Obernier, K., White, K. M., O’Meara, M. J., Rezelj, V. V., Guo, J. Z., Swaney, D. L., Tummino, T. A., Huttenhain, R., Kaake, R. M., Richards, A. L., Tutuncuoglu, B., Foussard, H., Batra, J., Haas, K., Modak, M., Kim, M., Haas, P., Polacco, B. J., Braberg, H., Fabius, J. M., Eckhardt, M., Soucheray, M., Bennett, M. J., Cakir, M., McGregor, M. J., Li, Q., Meyer, B., Roesch, F., Vallet, T., Mac Kain, A., Miorin, L., Moreno, E., Naing, Z. Z. C., Zhou, Y., Peng, S., Shi, Y., Zhang, Z., Shen, W., Kirby, I. T., Melnyk, J. E., Chorba, J. S., Lou, K., Dai, S. A., Barrio-Hernandez, I., Memon, D., Hernandez-Armenta, C., Lyu, J., Mathy, C. J. P., Perica, T., Pilla, K. B., Ganesan, S. J., Saltzberg, D. J., Rakesh, R., Liu, X., Rosenthal, S. B., Calviello, L., Venkataramanan, S., Liboy-Lugo, J., Lin, Y., Huang, X. P., Liu, Y., Wankowicz, S. A., Bohn, M., Safari, M., Ugur, F. S., Koh, C., Savar, N. S., Tran, Q. D., Shengjuler, D., Fletcher, S. J., O’Neal, M. C., Cai, Y., Chang, J. C. J., Broadhurst, D. J., Klippsten, S., Sharp, P. P., Wenzell, N. A., Kuzuoglu-Ozturk, D., Wang, H. Y., Trenker, R., Young, J. M., Cavero, D. A., Hiatt, J., Roth, T. L., Rathore, U., Subramanian, A., Noack, J., Hubert, M., Stroud, R. M., Frankel, A. D., Rosenberg, O. S., Verba, K. A., Agard, D. A., Ott, M., Emerman, M., Jura, N., von Zastrow, M., Verdin, E., Ashworth, A., Schwartz, O., d’Enfert, C., Mukherjee, S., Jacobson, M., Malik, H. S., Fujimori, D. G., Ideker, T., Craik, C. S., Floor, S. N., Fraser, J. S., Gross, J. D., Sali, A., Roth, B. L., Ruggero, D., Taunton, J., Kortemme, T., Beltrao, P., Vignuzzi, M., Garcia-Sastre, A., Shokat, K. M., Shoichet, B. K., and Krogan, N. J. (2020) A SARS-CoV-2 protein interaction map reveals targets for drug repurposing. Nature 583, 459–468

14. Puthenveetil, R., Lun, C. M., Murphy, R. E., Healy, L. B., Vilmen, G., Christenson, E. T., Freed, E. O., and Banerjee, A. (2021) S-acylation of SARS-CoV-2 Spike Protein: Mechanistic Dissection, In Vitro Reconstitution and Role in Viral Infectivity. J Biol Chem, 101112

15. Wu, Z., Zhang, Z., Wang, X., Zhang, J., Ren, C., Li, Y., Gao, L., Liang, X., Wang, P., and Ma, C. (2021) Palmitoylation of SARS-CoV-2 S protein is essential for viral infectivity. Signal Transduct Target Ther 6, 231

16. Mesquita, F., Abrami, L., Sergeeva, O., Turelli, P., Kunz, B., Raclot, C., Montoya, J., Abriata, L., Peraro, M., Trono, D., D’Angelo, G., and van der Goot, F. (2021) S-acylation controls SARS-Cov-2 membrane lipid organization and enhances infectivity. BioRXIV

17. Mesquita, F. S., Abrami, L., Sergeeva, O., Turelli, P., Qing, E., Kunz, B., Raclot, C., Paz Montoya, J., Abriata, L. A., Gallagher, T., Dal Peraro, M., Trono, D., D’Angelo, G., and van der Goot, F. G. (2021) S-acylation controls SARS-CoV-2 membrane lipid organization and enhances infectivity. Dev Cell 56, 2790–2807 e2798

18. Zeng, X. T., Yu, X. X., and Cheng, W. (2021) The interactions of ZDHHC5/GOLGA7 with SARS-CoV-2 spike (S) protein and their effects on S protein’s subcellular localization, palmitoylation and pseudovirus entry. Virol J 18, 257

19. Li, D., Liu, Y., Lu, Y., Gao, S., and Zhang, L. (2022) Palmitoylation of SARS-CoV-2 S protein is critical for S-mediated syncytia formation and virus entry. J Med Virol 94, 342–348

20. Ramadan, A. A., Mayilsamy, K., McGill, A. R., Ghosh, A., Giulianotti, M. A., Donow, H. M., Mohapatra, S. S., Mohapatra, S., Chandran, B., Deschenes, R. J., and Roy, A. (2022) Identification of SARS-CoV-2 Spike Palmitoylation Inhibitors That Results in Release of Attenuated Virus with Reduced Infectivity. Viruses 14

21. Thorp, E. B., Boscarino, J. A., Logan, H. L., Goletz, J. T., and Gallagher, T. M. (2006) Palmitoylations on murine coronavirus spike proteins are essential for virion assembly and infectivity. J Virol 80, 1280–1289

22. Shulla, A., and Gallagher, T. (2009) Role of spike protein endodomains in regulating coronavirus entry. J Biol Chem 284, 32725–32734

23. Petit, C. M., Chouljenko, V. N., Iyer, A., Colgrove, R., Farzan, M., Knipe, D. M., and Kousoulas, K. G. (2007) Palmitoylation of the cysteine-rich endodomain of the SARS-coronavirus spike glycoprotein is important for spike-mediated cell fusion. Virology 360, 264–274

24. Molday, L. L., and Molday, R. S. (2014) 1D4: a versatile epitope tag for the purification and characterization of expressed membrane and soluble proteins. Methods Mol Biol 1177, 1–15

25. Shang, J., Ye, G., Shi, K., Wan, Y., Luo, C., Aihara, H., Geng, Q., Auerbach, A., and Li, F. (2020) Structural basis of receptor recognition by SARS-CoV-2. Nature 581, 221–224

26. Ou, X., Liu, Y., Lei, X., Li, P., Mi, D., Ren, L., Guo, L., Guo, R., Chen, T., Hu, J., Xiang, Z., Mu, Z., Chen, X., Chen, J., Hu, K., Jin, Q., Wang, J., and Qian, Z. (2020) Characterization of spike glycoprotein of SARS-CoV-2 on virus entry and its immune cross-reactivity with SARS-CoV. Nat Commun 11, 1620

27. Gaebler, A., Milan, R., Straub, L., Hoelper, D., Kuerschner, L., and Thiele, C. (2013) Alkyne lipids as substrates for click chemistry-based in vitro enzymatic assays. J Lipid Res 54, 2282–2290

28. Tornoe, C. W., Christensen, C., and Meldal, M. (2002) Peptidotriazoles on solid phase: [1,2,3]-triazoles by regiospecific copper(i)-catalyzed 1,3-dipolar cycloadditions of terminal alkynes to azides. J Org Chem 67, 3057–3064

29. Rostovtsev, V. V., Green, L. G., Fokin, V. V., and Sharpless, K. B. (2002) A stepwise huisgen cycloaddition process: copper(I)-catalyzed regioselective “ligation” of azides and terminal alkynes. Angew Chem Int Ed Engl 41, 2596–2599

30. Charron, G., Zhang, M. M., Yount, J. S., Wilson, J., Raghavan, A. S., Shamir, E., and Hang, H. C. (2009) Robust fluorescent detection of protein fatty-acylation with chemical reporters. J Am Chem Soc 131, 4967–4975

31. Percher, A., Ramakrishnan, S., Thinon, E., Yuan, X., Yount, J. S., and Hang, H. C. (2016) Mass-tag labeling reveals site-specific and endogenous levels of protein S-fatty acylation. Proc Natl Acad Sci U S A 113, 4302–4307

32. Forrester, M. T., Hess, D. T., Thompson, J. W., Hultman, R., Moseley, M. A., Stamler, J. S., and Casey, P. J. (2011) Site-specific analysis of protein S-acylation by resin-assisted capture. J Lipid Res 52, 393–398

33. Abe, K. T., Li, Z., Samson, R., Samavarchi-Tehrani, P., Valcourt, E. J., Wood, H., Budylowski, P., Dupuis, A. P., 2nd, Girardin, R. C., Rathod, B., Wang, J. H., Barrios-Rodiles, M., Colwill, K., McGeer, A. J., Mubareka, S., Gommerman, J. L., Durocher, Y., Ostrowski, M., McDonough, K. A., Drebot, M. A., Drews, S. J., Rini, J. M., and Gingras, A. C. (2020) A simple protein-based surrogate neutralization assay for SARS-CoV-2. JCI Insight 5

34. Bosch, B. J., van der Zee, R., de Haan, C. A., and Rottier, P. J. (2003) The coronavirus spike protein is a class I virus fusion protein: structural and functional characterization of the fusion core complex. J Virol 77, 8801–8811

35. Cattin-Ortola, J., Welch, L. G., Maslen, S. L., Papa, G., James, L. C., and Munro, S. (2021) Sequences in the cytoplasmic tail of SARS-CoV-2 Spike facilitate expression at the cell surface and syncytia formation. Nat Commun 12, 5333

36. Jennings, B. C., and Linder, M. E. (2012) DHHC protein S-acyltransferases use similar ping-pong kinetic mechanisms but display different acyl-CoA specificities. J Biol Chem 287, 7236–7245

37. Nomura, R., Kiyota, A., Suzaki, E., Kataoka, K., Ohe, Y., Miyamoto, K., Senda, T., and Fujimoto, T. (2004) Human coronavirus 229E binds to CD13 in rafts and enters the cell through caveolae. J Virol 78, 8701–8708

38. Fekri, F., Abousawan, J., Bautista, S., Orofiamma, L., Dayam, R. M., Antonescu, C. N., and Karshafian, R. (2019) Targeted enhancement of flotillin-dependent endocytosis augments cellular uptake and impact of cytotoxic drugs. Sci Rep 9, 17768

39. Li, Z., Tomlinson, A. C., Wong, A. H., Zhou, D., Desforges, M., Talbot, P. J., Benlekbir, S., Rubinstein, J. L., and Rini, J. M. (2019) The human coronavirus HCoV-229E S-protein structure and receptor binding. Elife 8

40. Lu, Y., Zheng, Y., Coyaud, E., Zhang, C., Selvabaskaran, A., Yu, Y., Xu, Z., Weng, X., Chen, J. S., Meng, Y., Warner, N., Cheng, X., Liu, Y., Yao, B., Hu, H., Xia, Z., Muise, A. M., Klip, A., Brumell, J. H., Girardin, S. E., Ying, S., Fairn, G. D., Raught, B., Sun, Q., and Neculai, D. (2019) Palmitoylation of NOD1 and NOD2 is required for bacterial sensing. Science 366, 460–467

41. Loftus, T. M., Jaworsky, D. E., Frehywot, G. L., Townsend, C. A., Ronnett, G. V., Lane, M. D., and Kuhajda, F. P. (2000) Reduced food intake and body weight in mice treated with fatty acid synthase inhibitors. Science 288, 2379–2381

42. Montesdeoca, N., Lopez, M., Ariza, X., Herrero, L., and Makowski, K. (2020) Inhibitors of lipogenic enzymes as a potential therapy against cancer. FASEB J 34, 11355–11381

43. Flavin, R., Peluso, S., Nguyen, P. L., and Loda, M. (2010) Fatty acid synthase as a potential therapeutic target in cancer. Future Oncol 6, 551–562

44. Singh, S., Karthikeyan, C., and Moorthy, N. (2020) Recent Advances in the Development of Fatty Acid Synthase Inhibitors as Anticancer Agents. Mini Rev Med Chem 20, 1820–1837

45. Syed-Abdul, M. M., Parks, E. J., Gaballah, A. H., Bingham, K., Hammoud, G. M., Kemble, G., Buckley, D., McCulloch, W., and Manrique-Acevedo, C. (2020) Fatty Acid Synthase Inhibitor TVB-2640 Reduces Hepatic de Novo Lipogenesis in Males With Metabolic Abnormalities. Hepatology 72, 103–118

46. Heuer, T. S., Ventura, R., Mordec, K., Lai, J., Fridlib, M., Buckley, D., and Kemble, G. (2017) FASN Inhibition and Taxane Treatment Combine to Enhance Anti-tumor Efficacy in Diverse Xenograft Tumor Models through Disruption of Tubulin Palmitoylation and Microtubule Organization and FASN Inhibition-Mediated Effects on Oncogenic Signaling and Gene Expression. EBioMedicine 16, 51–62

47. Ventura, R., Mordec, K., Waszczuk, J., Wang, Z., Lai, J., Fridlib, M., Buckley, D., Kemble, G., and Heuer, T. S. (2015) Inhibition of de novo Palmitate Synthesis by Fatty Acid Synthase Induces Apoptosis in Tumor Cells by Remodeling Cell Membranes, Inhibiting Signaling Pathways, and Reprogramming Gene Expression. EBioMedicine 2, 808–824

48. Davda, D., El Azzouny, M. A., Tom, C. T., Hernandez, J. L., Majmudar, J. D., Kennedy, R. T., and Martin, B. R. (2013) Profiling targets of the irreversible palmitoylation inhibitor 2-bromopalmitate. ACS Chem Biol 8, 1912–1917

49. Banerjee, A., Nasir, J. A., Budylowski, P., Yip, L., Aftanas, P., Christie, N., Ghalami, A., Baid, K., Raphenya, A. R., Hirota, J. A., Miller, M. S., McGeer, A. J., Ostrowski, M., Kozak, R. A., McArthur, A. G., Mossman, K., and Mubareka, S. (2020) Isolation, Sequence, Infectivity, and Replication Kinetics of Severe Acute Respiratory Syndrome Coronavirus 2. Emerg Infect Dis 26, 2054–2063

50. Loomba, R., Mohseni, R., Lucas, K. J., Gutierrez, J. A., Perry, R. G., Trotter, J. F., Rahimi, R. S., Harrison, S. A., Ajmera, V., Wayne, J. D., O’Farrell, M., McCulloch, W., Grimmer, K., Rinella, M., Wai-Sun Wong, V., Ratziu, V., Gores, G. J., Neuschwander-Tetri, B. A., and Kemble, G. (2021) TVB-2640 (FASN Inhibitor) for the Treatment of Nonalcoholic Steatohepatitis: FASCINATE-1, a Randomized, Placebo-Controlled Phase 2a Trial. Gastroenterology 161, 1475–1486

51. Falchook, G., Infante, J., Arkenau, H. T., Patel, M. R., Dean, E., Borazanci, E., Brenner, A., Cook, N., Lopez, J., Pant, S., Frankel, A., Schmid, P., Moore, K., McCulloch, W., Grimmer, K., O’Farrell, M., Kemble, G., and Burris, H. (2021) First-in-human study of the safety, pharmacokinetics, and pharmacodynamics of first-in-class fatty acid synthase inhibitor TVB-2640 alone and with a taxane in advanced tumors. EClinicalMedicine 34, 100797

52. Ohol, Y. M., Wang, Z., Kemble, G., and Duke, G. (2015) Direct Inhibition of Cellular Fatty Acid Synthase Impairs Replication of Respiratory Syncytial Virus and Other Respiratory Viruses. PLoS One 10, e0144648

53. Kim, Y. C., Lee, S. E., Kim, S. K., Jang, H. D., Hwang, I., Jin, S., Hong, E. B., Jang, K. S., and Kim, H. S. (2019) Toll-like receptor mediated inflammation requires FASN-dependent MYD88 palmitoylation. Nat Chem Biol 15, 907–916

54. Jochen, A. L., Hays, J., and Mick, G. (1995) Inhibitory effects of cerulenin on protein palmitoylation and insulin internalization in rat adipocytes. Biochim Biophys Acta 1259, 65–72

55. Xiong, W., Sun, K. Y., Zhu, Y., Zhang, X., Zhou, Y. H., and Zou, X. (2021) Metformin alleviates inflammation through suppressing FASN-dependent palmitoylation of Akt. Cell Death Dis 12, 934

56. Wei, X., Schneider, J. G., Shenouda, S. M., Lee, A., Towler, D. A., Chakravarthy, M. V., Vita, J. A., and Semenkovich, C. F. (2011) De novo lipogenesis maintains vascular homeostasis through endothelial nitric-oxide synthase (eNOS) palmitoylation. J Biol Chem 286, 2933–2945

57. Wei, X., Yang, Z., Rey, F. E., Ridaura, V. K., Davidson, N. O., Gordon, J. I., and Semenkovich, C. F. (2012) Fatty acid synthase modulates intestinal barrier function through palmitoylation of mucin 2. Cell Host Microbe 11, 140–152

58. Weis, M. T., Crumley, J. L., Young, L. H., and Stallone, J. N. (2004) Inhibiting long chain fatty Acyl CoA synthetase increases basal and agonist-stimulated NO synthesis in endothelium. Cardiovasc Res 63, 338–346

59. Kilian, N., Zhang, Y., LaMonica, L., Hooker, G., Toomre, D., Mamoun, C. B., and Ernst, A. M. (2020) Palmitoylated Proteins in Plasmodium falciparum-Infected Erythrocytes: Investigation with Click Chemistry and Metabolic Labeling. Bioessays 42, e1900145

60. Leonardi, R., Zhang, Y. M., Rock, C. O., and Jackowski, S. (2005) Coenzyme A: back in action. Prog Lipid Res 44, 125–153

61. Karthigeyan, K. P., Zhang, L., Loiselle, D. R., Haystead, T. A. J., Bhat, M., Yount, J. S., and Kwiek, J. J. (2021) A bioorthogonal chemical reporter for fatty acid synthase-dependent protein acylation. J Biol Chem 297, 101272

62. Topolska, M., Martinez-Montanes, F., and Ejsing, C. S. (2020) A Simple and Direct Assay for Monitoring Fatty Acid Synthase Activity and Product-Specificity by High-Resolution Mass Spectrometry. Biomolecules 10

63. Boscarino, J. A., Logan, H. L., Lacny, J. J., and Gallagher, T. M. (2008) Envelope protein palmitoylations are crucial for murine coronavirus assembly. J Virol 82, 2989–2999

64. St-Germain, J. R., Astori, A., and Raught, B. (2021) A SARS-CoV-2 Peptide Spectral Library Enables Rapid, Sensitive Identification of Virus Peptides in Complex Biological Samples. J Proteome Res 20, 2187–2194

65. Dixon, C. L., Mekhail, K., and Fairn, G. D. (2021) Examining the Underappreciated Role of S-Acylated Proteins as Critical Regulators of Phagocytosis and Phagosome Maturation in Macrophages. Front Immunol 12, 659533

66. Kang, Y. L., Chou, Y. Y., Rothlauf, P. W., Liu, Z., Soh, T. K., Cureton, D., Case, J. B., Chen, R. E., Diamond, M. S., Whelan, S. P. J., and Kirchhausen, T. (2020) Inhibition of PIKfyve kinase prevents infection by Zaire ebolavirus and SARS-CoV-2. Proc Natl Acad Sci U S A 117, 20803–20813

67. Williams, C. G., Jureka, A. S., Silvas, J. A., Nicolini, A. M., Chvatal, S. A., Carlson-Stevermer, J., Oki, J., Holden, K., and Basler, C. F. (2021) Inhibitors of VPS34 and fatty-acid metabolism suppress SARS-CoV-2 replication. Cell Rep 36, 109479

68. Sauvat, A., Ciccosanti, F., Colavita, F., Di Rienzo, M., Castilletti, C., Capobianchi, M. R., Kepp, O., Zitvogel, L., Fimia, G. M., Piacentini, M., and Kroemer, G. (2020) On-target versus off-target effects of drugs inhibiting the replication of SARS-CoV-2. Cell Death Dis 11, 656

69. Shen, B., Yi, X., Sun, Y., Bi, X., Du, J., Zhang, C., Quan, S., Zhang, F., Sun, R., Qian, L., Ge, W., Liu, W., Liang, S., Chen, H., Zhang, Y., Li, J., Xu, J., He, Z., Chen, B., Wang, J., Yan, H., Zheng, Y., Wang, D., Zhu, J., Kong, Z., Kang, Z., Liang, X., Ding, X., Ruan, G., Xiang, N., Cai, X., Gao, H., Li, L., Li, S., Xiao, Q., Lu, T., Zhu, Y., Liu, H., Chen, H., and Guo, T. (2020) Proteomic and Metabolomic Characterization of COVID-19 Patient Sera. Cell 182, 59–72 e15

70. Chu, J., Xing, C., Du, Y., Duan, T., Liu, S., Zhang, P., Cheng, C., Henley, J., Liu, X., Qian, C., Yin, B., Wang, H. Y., and Wang, R. F. (2021) Pharmacological inhibition of fatty acid synthesis blocks SARS-CoV-2 replication. Nat Metab

71. Dias, S. S. G., Soares, V. C., Ferreira, A. C., Sacramento, C. Q., Fintelman-Rodrigues, N., Temerozo, J. R., Teixeira, L., Nunes da Silva, M. A., Barreto, E., Mattos, M., de Freitas, C. S., Azevedo-Quintanilha, I. G., Manso, P. P. A., Miranda, M. D., Siqueira, M. M., Hottz, E. D., Pao, C. R. R., Bou-Habib, D. C., Barreto-Vieira, D. F., Bozza, F. A., Souza, T. M. L., and Bozza, P. T. (2020) Lipid droplets fuel SARS-CoV-2 replication and production of inflammatory mediators. PLoS Pathog 16, e1009127

72. Toelzer, C., Gupta, K., Yadav, S. K. N., Borucu, U., Davidson, A. D., Kavanagh Williamson, M., Shoemark, D. K., Garzoni, F., Staufer, O., Milligan, R., Capin, J., Mulholland, A. J., Spatz, J., Fitzgerald, D., Berger, I., and Schaffitzel, C. (2020) Free fatty acid binding pocket in the locked structure of SARS-CoV-2 spike protein. Science 370, 725–730

73. Jones, S. F., and Infante, J. R. (2015) Molecular Pathways: Fatty Acid Synthase. Clin Cancer Res 21, 5434–5438

74. Korbie, D. J., and Mattick, J. S. (2008) Touchdown PCR for increased specificity and sensitivity in PCR amplification. Nat Protoc 3, 1452–1456

75. Milewska, A., Kaminski, K., Ciejka, J., Kosowicz, K., Zeglen, S., Wojarski, J., Nowakowska, M., Szczubialka, K., and Pyrc, K. (2016) HTCC: Broad Range Inhibitor of Coronavirus Entry. PLoS One 11, e0156552

76. Baer, A., and Kehn-Hall, K. (2014) Viral concentration determination through plaque assays: using traditional and novel overlay systems. J Vis Exp, e52065

77. De Albuquerque, N., Baig, E., Ma, X., Zhang, J., He, W., Rowe, A., Habal, M., Liu, M., Shalev, I., Downey, G. P., Gorczynski, R., Butany, J., Leibowitz, J., Weiss, S. R., McGilvray, I. D., Phillips, M. J., Fish, E. N., and Levy, G. A. (2006) Murine hepatitis virus strain 1 produces a clinically relevant model of severe acute respiratory syndrome in A/J mice. J Virol 80, 10382–10394

78. Sugiyama, M. G., Armstrong, S. M., Wang, C., Hwang, D., Leong-Poi, H., Advani, A., Advani, S., Zhang, H., Szaszi, K., Tabuchi, A., Kuebler, W. M., Van Slyke, P., Dumont, D. J., and Lee, W. L. (2015) The Tie2-agonist Vasculotide rescues mice from influenza virus infection. Sci Rep 5, 11030

79. Choi, H., Larsen, B., Lin, Z. Y., Breitkreutz, A., Mellacheruvu, D., Fermin, D., Qin, Z. S., Tyers, M., Gingras, A. C., and Nesvizhskii, A. I. (2011) SAINT: probabilistic scoring of affinity purification-mass spectrometry data. Nat Methods 8, 70–73

